# A unified law for inhibitory control in active dendrites

**DOI:** 10.64898/2026.05.15.725398

**Authors:** Yutao He, Gan He, Boxiang Huang, Panayiota Poirazi, Tiejun Huang, Kai Du

## Abstract

Neuronal computation depends on the balance between excitation and inhibition, yet how this balance is implemented across the dendritic tree remains unclear. Classical views predict that inhibition should be most effective near the soma or along the path from excitation to output, but many interneuron subtypes preferentially target remote dendritic compartments. This apparent paradox is sharpened by active dendrites, where local NMDA spikes, calcium plateaus and backpropagating action potentials can make distal branches powerful contributors to somatic firing. Here we develop an analytical framework that extracts general principles of inhibition from biophysically detailed multi-compartment simulations. By reformulating the implicit voltage update of detailed neuron models as a matrix recursion, we derive exact voltage sensitivities to inhibitory synaptic perturbations. This leads to a unified Φ-a law: the somatic impact of inhibition factorizes into a global dendritic susceptibility term and a local synaptic perturbation term. Using this law to map inhibitory leverage and identify optimal inhibitory interventions, we show that active dendritic excitation can shift inhibitory hot zones from perisomatic regions toward distal or intermediate compartments. Across neocortical, hippocampal and striatal neuron models, the same response law explains convergent inhibitory strategies despite distinct cellular mechanisms. Our framework turns detailed numerical simulation into analytical theory, providing a general principle for how diverse dendritic inhibition controls active neurons.

## Introduction

The balance between excitation and inhibition (‘E-I balance’) is a fundamental requirement for stable computation in neural circuits^1-3^. Yet how this balance is implemented across the complex dendritic architecture of single neurons poses a central biophysical paradox. For decades, the logic of dendritic inhibition has been interpreted through the lens of static anatomy and passive cable theory^4^, according to which electrical signals attenuate with distance^5, 6^. Within this classical framework, inhibitory synapses are expected to be most effective when positioned proximally, acting primarily as local shunts or somatic counterweights to excitation^7^. Mammalian brains, however, have evolved a radically different architecture: inhibitory interneurons are extraordinarily diverse, their populations are often sparse^8, 9^, and many subtypes preferentially target electrically remote dendritic compartments rather than the perisomatic domain^10^ (Fig. 1a). Why a metabolically constrained biological system would preserve such apparently inefficient distal inhibition remains a central question.

**Fig. 1.**
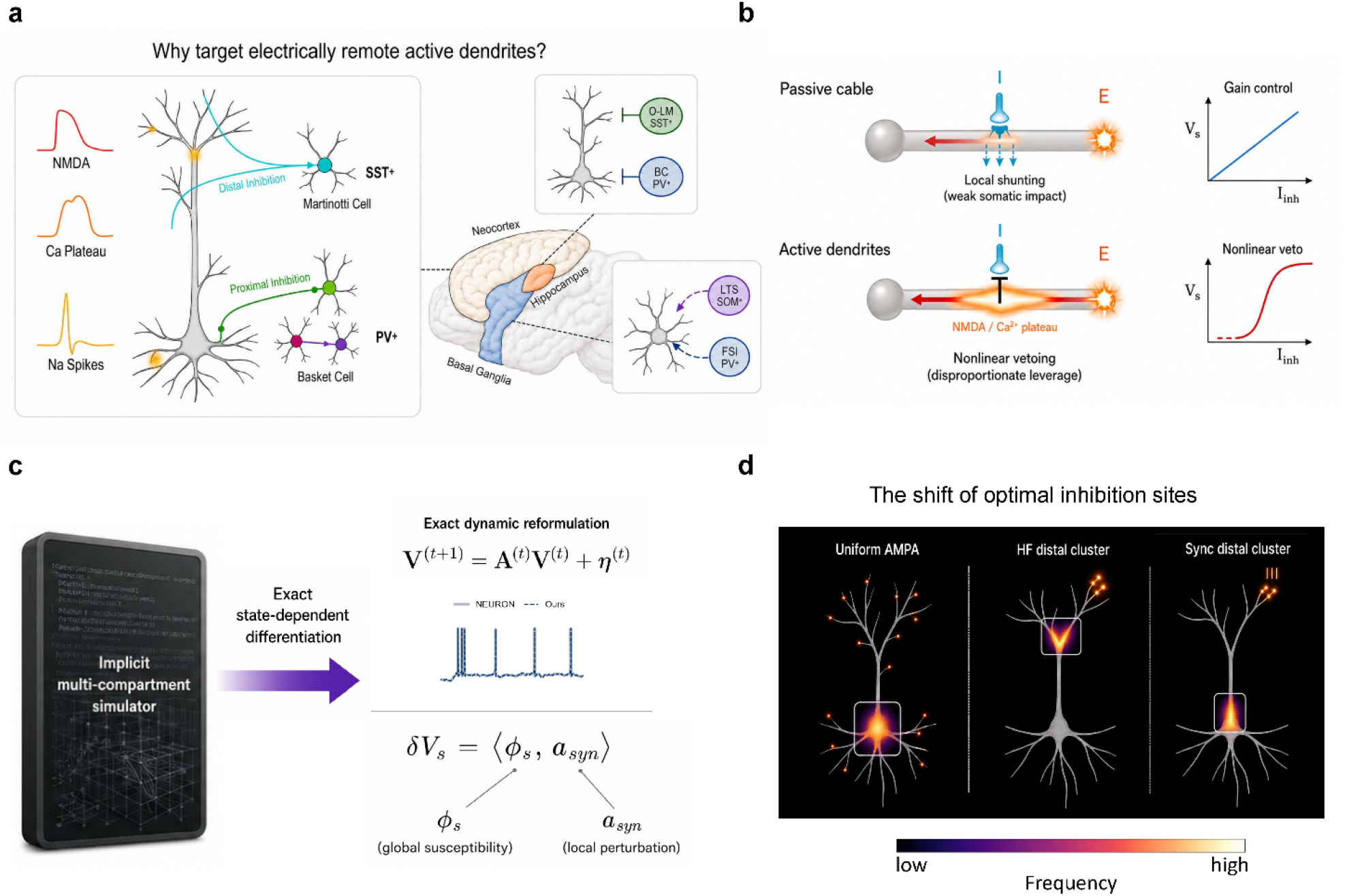
A state-dependent framework resolves the paradox of distal dendritic inhibition. (a) The anatomical paradox of distal inhibition. Across diverse brain circuits (including neocortex, hippocampus and basal ganglia), specialized interneuron subtypes such as SST^+^ Martinotti, O-LM and LTS cells preferentially target electrically remote dendritic compartments^8–10,43,44,58^ that can generate active regenerative events, including NMDA spikes and Ca^2+^ plateaus^11,12^. This architecture challenges classical passive-cable views that emphasize somatic proximity, as in PV^+^ basket-cell inhibition, for strong inhibitory control^4–7^. (b) Active dendrites enable nonlinear vetoing. In classical passive cables (top), off-path distal inhibition acts mainly as a weak local shunt, providing linear gain control over somatic voltage^17–20^. In active dendrites (bottom), local inhibition can intercept regenerative events, giving remote synapses disproportionate vetoing leverage over somatic output^13,26–30^. (c) Extraction of a unified response law. The implicit numerical update of a detailed multi-compartment model is analytically reformulated into a trajectory-conditioned dynamical system; state-dependent differentiation yields a response law in which the first-order somatic impact of a synaptic perturbation factorizes into global spatiotemporal susceptibility (Φs) and local synaptic drive (*a*_*syn*_). (d) Excitation-dependent shifts of inhibitory hot zones. Under this response law, optimal inhibitory sites are not fixed by anatomy, but dynamically reroute from perisomatic regions under uniform AMPA excitation to distal branches under high-frequency clustered inputs (source suppression), and toward the intermediate apical trunk under synchronous distal inputs (choke-point interception).

A growing body of physiological evidence suggests that this paradox arises because traditional frameworks understate the computational complexity of biological neurons. Dendrites are not passive cables, but active excitable structures endowed with voltage-gated conductances capable of generating highly nonlinear local events, including NMDA spikes and calcium plateau potentials^11, 12^. Under active regimes, the classical assumption of passive electrotonic attenuation breaks down. As a result, a distal synapse that appears electrically weak during quiescent states may become disproportionately powerful once local dendritic nonlinearities are engaged^12, 13^. The principles of dendritic inhibition can therefore no longer be adequately captured by anatomical distance alone, by impedance-based descriptors that are themselves voltage-, frequency-, and conductance-state dependent^14-16^, or by one-dimensional f/I curves^17-20^. Addressing this problem requires an analytical framework that can directly quantify how inhibitory perturbations propagate through active dendritic trees to shape somatic output.

Here we develop such a framework by reformulating the numerical update steps of detailed multi-compartment neuron models into an analytically tractable, differentiable dynamical system (Fig. 1c). This allows us to compute exact voltage gradients with respect to inhibitory synaptic parameters and thereby move from descriptive simulation to quantitative control analysis. On this basis, we define the optimal dendritic inhibition (ODI) problem: given an excitatory drive that elicits somatic spiking, what is the minimal inhibitory intervention—in strength, location, and timing—required to suppress it? Although the resulting spatiotemporal search space is combinatorially large, the analytical gradients extracted from our framework render its structure tractable.

Applying this framework across four canonical detailed neuron models—neocortical L2/3^21^ and L5 pyramidal cells^22^, hippocampal CA1 pyramidal neurons^23, 24^, and striatal medium spiny neurons^25^—reveals a remarkable principle of convergence under constraint. As structured excitation recruits local regenerative events, these architecturally diverse neuron classes converge on closely related macroscopic strategies of inhibitory deployment: the inhibitory “hot zone” shifts away from proximal regions toward the dendritic compartments engaged by the computation (Fig. 1d). Thus, the most effective site of inhibition is not fixed by static anatomy, but reorganizes with the state and spatial structure of excitation.

Analysis of the resulting gradients further reveals a unified response law for inhibitory control. We find that the somatic impact δV_s_ from a unit inhibitory perturbation can be factorized as

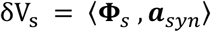

where **Φ**_s_ is a state-dependent susceptibility term determined by the neuron’s global biophysical state, and ***a***_*syn*_ is the local response to unit single-synapse perturbation. This decomposition shows that inhibitory efficacy is jointly set by global susceptibility and local drive. In neurons with strongly compartmentalized active dendrites, reconfiguration of the susceptibility term dominates shifts in inhibitory leverage. In more electrically compact neurons, by contrast, intrinsic susceptibility remains comparatively rigid, and state-dependent nonlinearities in the local perturbation term play a larger role. **Distinct neuron**

### classes therefore achieve macroscopic functional convergence through divergent biophysical mechanisms—an elegant manifestation of biological degeneracy

Together, this unified framework reconciles apparently conflicting views of dendritic inhibition by placing location-dependent, arithmetic and active-state mechanisms within a single response law ^13, 26-30^. By demonstrating that diverse neuron classes rely on the same fundamental response law—even if they exploit different terms within its decomposition—we provide a principled, quantitative rationale for the spatiotemporal diversity of dendritic inhibition. Rather than simply tiling a static anatomical tree, specialized interneuron subtypes dynamically tile a shifting landscape of inhibitory leverage.

## Results

### Active dendrites transform inhibition into a state-dependent control problem

Dendritic inhibition poses a state-dependent control problem for neurons. Inhibitory inputs are selectively deployed to regulate excitatory drive distributed across an extended dendritic tree, yet the efficacy of any single inhibitory event depends on the current biophysical state of that tree. Classical cable intuition predicts that inhibitory impact should decay with electrotonic distance and should therefore be strongest near the soma or along the direct path from excitation to output. Active dendrites complicate this picture: NMDA spikes, Ca^2+^ plateaus, and voltage-gated conductances can transiently turn electrically remote branches into strong determinants of somatic firing^6,11,12^ (Fig. 1b). The key question is therefore not only where inhibitory synapses are located anatomically, but where and when inhibition has maximal leverage over the current dendritic state.

Answering this question requires tracking how local synaptic perturbations propagate through realistic active dendrites. Detailed multi-compartment models contain the relevant morphology, channel distributions, and synaptic biophysics, but they are often used as forward simulators^22, 31, 32^: one specifies an input pattern, runs the model, and observes the voltage response. This makes it difficult to extract general laws of inhibitory control. We therefore took a different route. Rather than simplifying the neuron to obtain an analytical theory, we made the numerical simulator itself the object of analysis.

We reformulated the implicit voltage update used in detailed compartmental simulations into a trajectory-conditioned affine recursion:

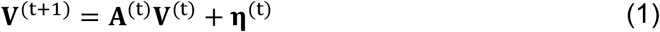

Here, **V**^(t)^ denotes the vector of membrane voltages across all compartments. The operator **A**^(t)^ and offset **η**^(t)^ are evaluated along the ongoing trajectory and encode the state-dependent contributions of passive, synaptic, and voltage-gated mechanisms. This expression is exact for the discretized simulator update rule. It does not remove the nonlinear biophysics, nor does it replace the neuron with a passive or globally linear approximation. Instead, it organizes the nonlinear numerical update into an explicit dynamical map.

Direct iteration of Eq. (1) reproduced the corresponding simulator trajectories with deviations limited to numerical tolerance and implementation-level differences (Fig. 2b and Supplementary Fig. 1). Thus, a realistic compartmental neuron can be treated not only as a numerical testbed, but also as an analyzable dynamical system. This simulator-level formulation provides a mathematical entry point into active dendrites while retaining the biophysical detail required for dendritic computation.

**Fig. 2.**
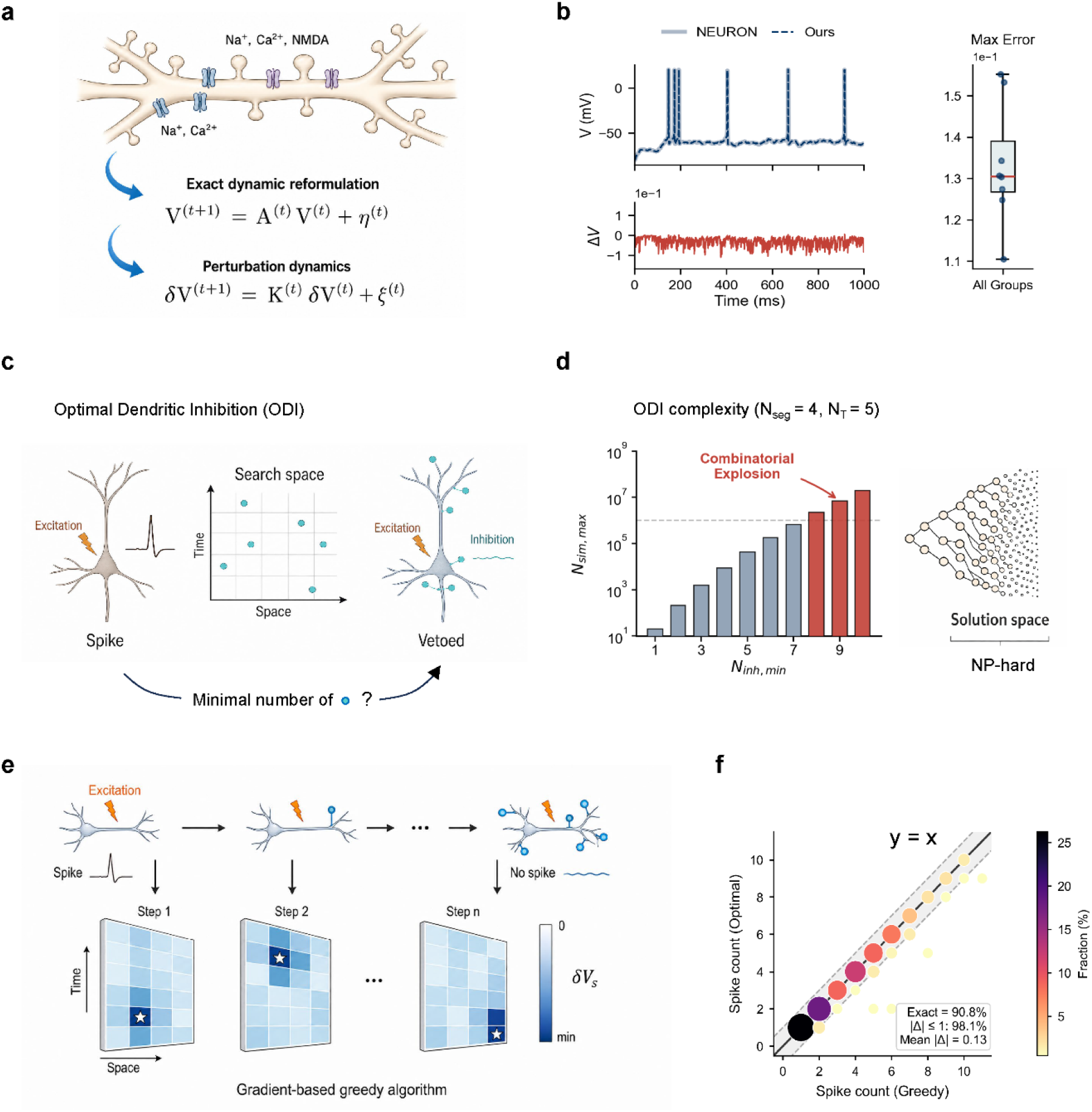
Simulator-derived sensitivities define optimal dendritic inhibition. (a) Exact dynamic reformulation of a detailed active dendrite. The implicit voltage update of a multi-compartment simulator is rewritten as a trajectory-conditioned affine recursion that preserves morphology, synaptic currents and voltage-dependent conductances; differentiating this recursion yields the first-order perturbation dynamics for inhibitory synaptic weights. (b) Direct iteration of the reformulated update reproduces voltage trajectories generated by NEURON^31,32^. Top, somatic voltage from the original simulator and the matrix computation; bottom, residual voltage error. The box plot summarizes maximum absolute error across simulation groups. (c) Schematic definition of the optimal dendritic inhibition (ODI) problem. Given an excitatory pattern that evokes a somatic spike, ODI seeks the minimal set of inhibitory events across dendritic space and time required to veto that spike. (d) Combinatorial complexity of ODI. The number of candidate inhibitory subsets increases rapidly with the minimal inhibitory count, yielding an NP-hard search over a high-dimensional solution space. (e) Gradient-guided greedy solution. At each iteration, candidate inhibitory events are ranked by their first-order somatic impact, the strongest vetoing event is selected, the trajectory is updated and sensitivities are recomputed until somatic spiking is suppressed. (f) Validation against exhaustive enumeration in a toy multi-compartment neuron. Points denote the frequency of greedy versus exact inhibitory spike counts; the dashed line marks equality. The greedy algorithm returns exact or near-exact solutions in most cases.

### Simulator-derived sensitivities define inhibitory leverage

Having obtained an explicit simulator-level dynamical map, we next asked how a candidate inhibitory synapse affects the voltage trajectory. Differentiating Eq. (1) with respect to the weight of a candidate inhibitory input yields an exact recursion for the first-order perturbation dynamics (Eq. (2); Methods). Here, **K**^(t)^ denotes the local Jacobian of the simulator-level map evaluated along the current trajectory, which propagates accumulated voltage perturbations across compartments and time. The source term ***ξ***^(t)^ denotes the direct perturbation induced by the candidate synaptic event. The resulting sensitivity quantifies the local derivative of somatic voltage with respect to an inhibitory event at a specific location and time, evaluated around the current trajectory.

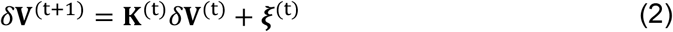

This sensitivity has a direct biological interpretation: it measures the inhibitory leverage of a candidate synaptic event under the current dendritic state. A high-leverage site is not necessarily the site closest to the soma, nor the site with the largest local IPSP. Rather, it is a spatiotemporal coordinate at which a local perturbation is efficiently transmitted through the active dendritic tree to influence somatic output. Thus, the gradient is not introduced merely as an optimization heuristic; it is the local first-order perturbation response of the active dendritic system itself.

These sensitivities allowed us to formulate the optimal dendritic inhibition (ODI) problem (Fig. 2c). Given an excitatory input pattern that elicits a somatic spike, we ask what is the smallest set of inhibitory events, within a specified candidate space of locations, timings, and synaptic weights, required to suppress that spike. This formulation captures a biologically relevant form of efficient inhibitory control: minimal inhibition that vetoes excitation. Because candidate inhibitory events span both dendritic space and time, the search space grows combinatorially, and the corresponding decision problem is NP-hard even under monotone-inhibition assumptions (Fig. 2d; Methods).

To obtain practical solutions in realistic models, we used a gradient-guided greedy strategy (Fig. 2e and Supplementary Fig. 2). At each step, the algorithm computes the inhibitory leverage of all candidate events, selects the event with the strongest negative first-order effect on somatic voltage, updates the neuron with that inhibitory event, and then recomputes the sensitivities around the newly updated trajectory. This repeated re-evaluation is essential. The method does not assume that inhibitory effects add linearly on a fixed background state; instead, it follows how the dendritic state itself changes as inhibition accumulates.

We validated this procedure in a simplified multi-compartment model for which exhaustive enumeration of the global optimum was still possible. The gradient-guided solutions closely approached the ground-truth optimum, typically requiring only a small number of additional inhibitory events relative to exhaustive search (Fig. 2f and Supplementary Fig. 2). Thus, simulator-derived sensitivities provide a practical route through the otherwise prohibitive search space of dendritic inhibition while preserving the nonlinear dynamics of the underlying model.

### A unified response law separates global dendritic susceptibility from local synaptic drive

Although the sensitivity recursion allows inhibitory leverage to be computed, it does not by itself explain why one inhibitory event is more effective than another. We therefore derived an explicit expression for the first-order somatic effect of a candidate inhibitory perturbation. For a candidate synapse *syn* and readout time t*, the first-order somatic response can be written as:

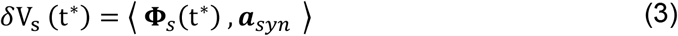

Here, ***a***_*syn*_ is the local voltage change induced by the unit perturbation on *syn*, whereas **Φ**_*s*_(t^∗^) is a trajectory-dependent spatiotemporal susceptibility kernel determined by the ongoing biophysical state of the neuron. For finite inhibitory perturbations, the actual somatic voltage change is approximated by this first-order term up to higher-order corrections.

This decomposition is the central theoretical result of the study. It shows that, to first order, inhibitory efficacy is governed by the alignment between two quantities (Fig. 3a). The first is local synaptic drive: how strongly an inhibitory input perturbs the membrane potential at its own site. The second is global dendritic susceptibility: how the current active dendritic state transforms that local perturbation into somatic impact. A large local IPSP is therefore not sufficient for strong somatic control; the perturbation must also occur at a location and time where the dendritic tree is susceptible to transmitting it to the soma.

**Fig. 3.**
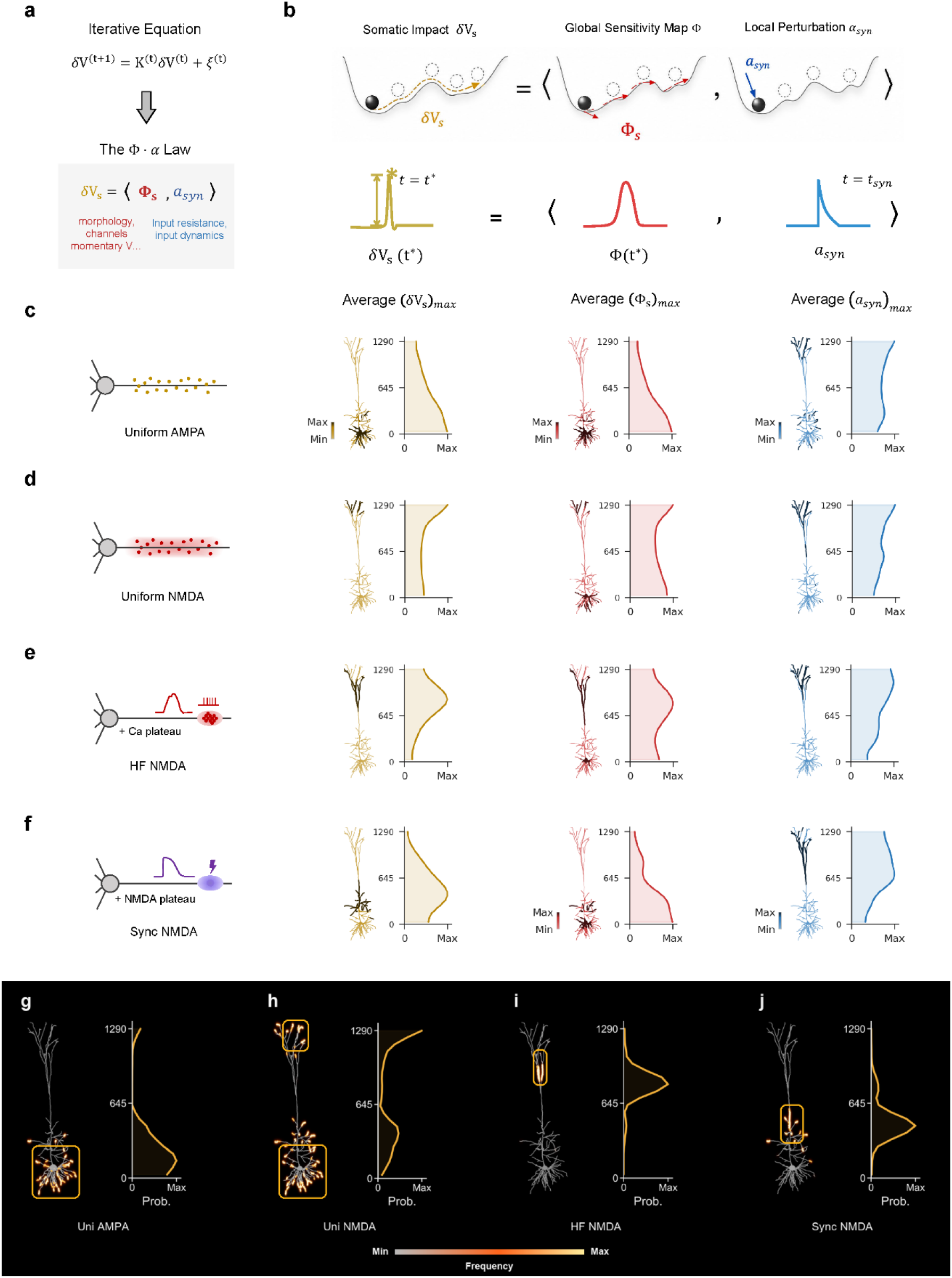
A Φ–a response law explains excitation-dependent spatial remapping of dendritic inhibition. (a) Differentiating the perturbation recursion yields the response law δVs = ⟨Φs, *a*_*syn*_⟩, separating the first-order somatic impact into a global susceptibility term (Φs) and a local synaptic perturbation term (*a*_*syn*_). (b) Schematic interpretation of the law. The somatic effect of an inhibitory event depends on the alignment between the state-dependent susceptibility landscape and the local synaptic drive, rather than on local IPSP amplitude or anatomical distance alone. (c–f) Excitation paradigms in a layer 5 pyramidal cell model^22^ and the corresponding averages of somatic impact, susceptibility and local perturbation: (c) uniform AMPA input; (d) uniform NMDA/AMPA input; (e) high-frequency distal NMDA/AMPA clusters that evoke a Ca^2+^ plateau; and (f) synchronous distal NMDA/AMPA clusters that evoke an NMDA plateau^11,33–37^. (g–j) ODI-derived spatial distributions of selected inhibitory events for the four regimes in c–f. Uniform AMPA favours basal/perisomatic control (g), uniform NMDA shifts inhibition outward (h), HF NMDA concentrates inhibition near the activated distal source (i), and Sync NMDA favours an intermediate apical choke point (j). Warm colours denote higher selection frequency or probability.

This response law connects experimental intuition with computational analysis. The local term ***a***_*j*_ corresponds to the immediate biophysical perturbation generated by synaptic current. The susceptibility kernel **Φ**_*s*_ corresponds to the state-dependent dendritic landscape through which this perturbation propagates (Fig. 3b). Active nonlinearities can reshape this landscape, creating transient conditions in which distal or intermediate compartments acquire unexpectedly high somatic leverage. Thus, Eq. (3) generalizes passive cable intuition: anatomical distance still matters, but its influence is dynamically gated by the nonlinear state of the dendritic tree.

Because both **Φ**_*s*_ and ***a***_*syn*_ are evaluated on the current trajectory, the response law is recomputed after the neuronal state changes. It is therefore a local law of active dendritic control, not a global linear superposition principle across arbitrary inhibitory patterns.

### Nonlinear dendritic events reroute optimal inhibition in space

We next used the ODI framework to ask how this response law operates under different excitation regimes. In a detailed layer 5 pyramidal neuron model ^22^, we compared four spatiotemporal patterns of excitation (Fig. 3c–f, left; Methods): uniform AMPA input (Uni AMPA), uniform NMDA/AMPA input (Uni NMDA), high-frequency distal NMDA/AMPA clusters (HF NMDA), and synchronous distal NMDA/AMPA clusters (Sync NMDA). These regimes were designed as idealized probes of experimentally established modes of pyramidal-cell excitation: sparse, distributed or asynchronous excitatory drive on the one hand, and clustered or temporally coordinated distal glutamatergic input capable of recruiting NMDA spikes and Ca^2+^ plateau potentials on the other. These patterns therefore span a progression from weakly structured distributed excitation to strongly nonlinear distal events^11,33-37^.

Under uniform AMPA input, dendritic integration remained comparatively weak and distributed. Distal branches were only modestly recruited, and the spatial projection of the susceptibility kernel decreased broadly from proximal to distal compartments. Consistent with this regime, optimal inhibition was concentrated mainly in perisomatic and basal compartments (Fig. 3c,g).

Adding distributed NMDA/AMPA input increased overall dendritic excitability and partially shifted the inhibitory hot zone outward (Fig. 3d,h). This indicates that even broadly distributed nonlinear conductance can alter the leverage landscape and allow more distal compartments to contribute to efficient inhibitory control.

The two structured distal regimes produced more pronounced state-dependent remapping. Under HF NMDA, repetitive clustered input generated a sustained local Ca^2+^ plateau in the stimulated distal branch. In this condition, elevated local drive and enhanced distal susceptibility were associated with a shift of the optimal inhibitory hot zone toward the activated distal branch (Fig. 3e,i). The ODI solution therefore implements a source-suppression strategy: inhibition is most effective when placed near the branch generating the sustained nonlinear event.

By contrast, Sync NMDA also generated a strong distal nonlinear event, but the optimal hot zone shifted toward the intermediate apical trunk rather than remaining concentrated in the distal tuft (Fig. 3j). In this condition, the local perturbation term remained large near the distal input, but the susceptibility landscape was preferentially elevated along intermediate compartments (Fig. 3f). This pattern is consistent with a choke-point mechanism, in which inhibition most effectively controls somatic output by intercepting the influence of a synchronous distal event along its route to the soma. Thus, superficially similar distal nonlinear events can require different inhibitory strategies because they reshape the local drive and global susceptibility terms in different ways.

### Susceptibility dynamics determine the timing of inhibition

Efficient inhibition also reorganized in time. Under HF NMDA, optimal inhibitory events were distributed broadly across the stimulation period. Under Sync NMDA, optimal inhibition was concentrated into more discrete temporal windows. Quantification with a Gini coefficient confirmed that Sync NMDA produced more temporally uneven inhibition, whereas HF NMDA produced more temporally dispersed inhibition (Fig. 4f).

**Fig. 4.**
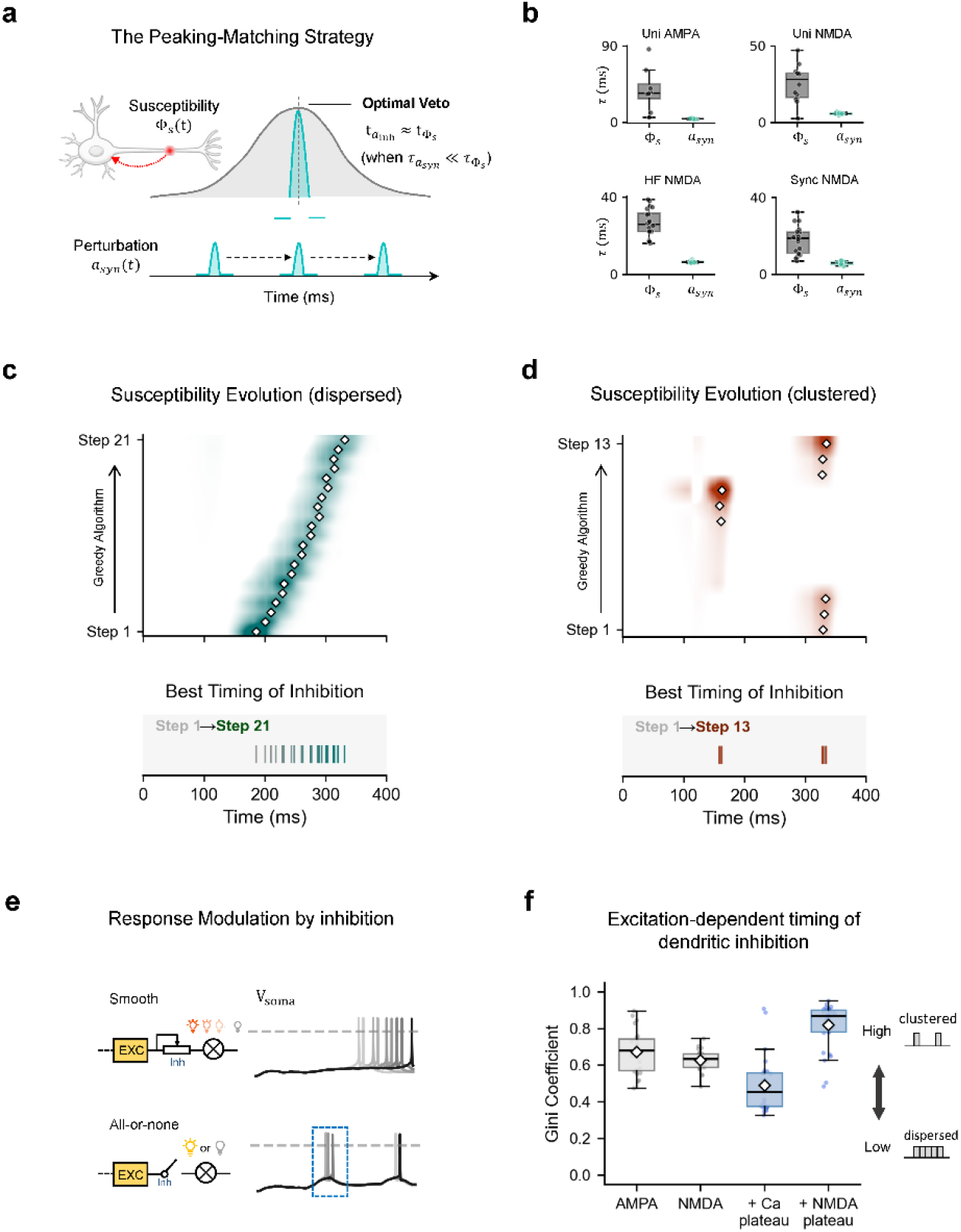
Susceptibility dynamics set the timing of optimal inhibition. (a) Peaking-matching strategy. A brief GABAergic perturbation is maximally effective when its local waveform is aligned with a peak in the susceptibility kernel, so the optimal inhibitory latency is determined by the temporal structure of Φs. (b) Characteristic time scales of susceptibility and local perturbation under four excitation regimes. Susceptibility evolves on broader, state-dependent time scales than the local synaptic waveform. (c,d) Evolution of susceptibility peaks during greedy inhibition for dispersed (HF NMDA; c) and clustered (Sync NMDA; d) regimes. White markers denote selected inhibitory events, and lower rasters show the best inhibitory timing at each algorithmic step. (e) Schematic somatic responses illustrating smooth versus all-or-none modulation by inhibition. Smooth regimes shift spike timing gradually, whereas all-or-none regimes switch between discrete somatic responses. (f) Quantification of temporal clustering using the Gini coefficient. Higher values indicate more clustered inhibitory timing. Synchronous plateau-evoking input produces more temporally concentrated inhibition than high-frequency Ca^2+^-plateau input.

This temporal organization follows from the same response law. Fast GABAergic perturbations have brief local waveforms, whereas the susceptibility kernel evolves over a broader timescale determined by the dendritic state (Fig. 4b). Inhibitory efficacy is therefore maximized when the brief synaptic perturbation is temporally aligned with peaks of ϕ_s_ (Fig. 4a). The timing of optimal inhibition is not an independent feature of the algorithm; it reflects the temporal geometry of the susceptibility kernel.

Under HF NMDA, susceptibility peaks shifted smoothly over time as inhibition accumulated, producing distributed inhibitory timings (Fig. 4c). Under Sync NMDA, susceptibility peaks became concentrated in discrete temporal windows, producing clustered inhibition (Fig. 4d). These dynamics were mirrored in the somatic response: under HF NMDA, inhibition gradually shifted the timing of the first somatic spike, whereas under Sync NMDA, spike timing switched more abruptly between discrete states. Thus, different excitation regimes impose different temporal structures on dendritic susceptibility, and optimal inhibition follows those structures.

### Distinct neuron classes implement a shared inhibitory logic through different mechanisms

Finally, we asked whether these principles generalize beyond the layer 5 pyramidal neuron. We applied the same framework to three additional detailed neuron models with distinct morphologies and electrotonic structures: a layer 2/3 pyramidal neuron^21^, a hippocampal CA1 pyramidal neuron^23^, and a striatal medium spiny neuron^25^.

Across the models examined, excitation regime produced a shared qualitative reorganization of inhibitory control (Fig. 5a). Weak distributed excitation favored somatically adjacent or basal/proximal compartments. High-frequency clustered excitation shifted inhibition toward the stimulated distal dendritic region. Synchronous clustered excitation shifted inhibition back toward intermediate compartments or the direct route from the stimulated region to the soma. Thus, across these morphologically distinct models, optimal inhibition followed a conserved outward-then-retracted spatial trajectory rather than a monotonic rule based on anatomical distance alone (Fig. 5b). Temporal organization was also conserved: synchronous distal excitation produced more clustered inhibition than high-frequency distal excitation (Fig. 5c).

**Fig. 5.**
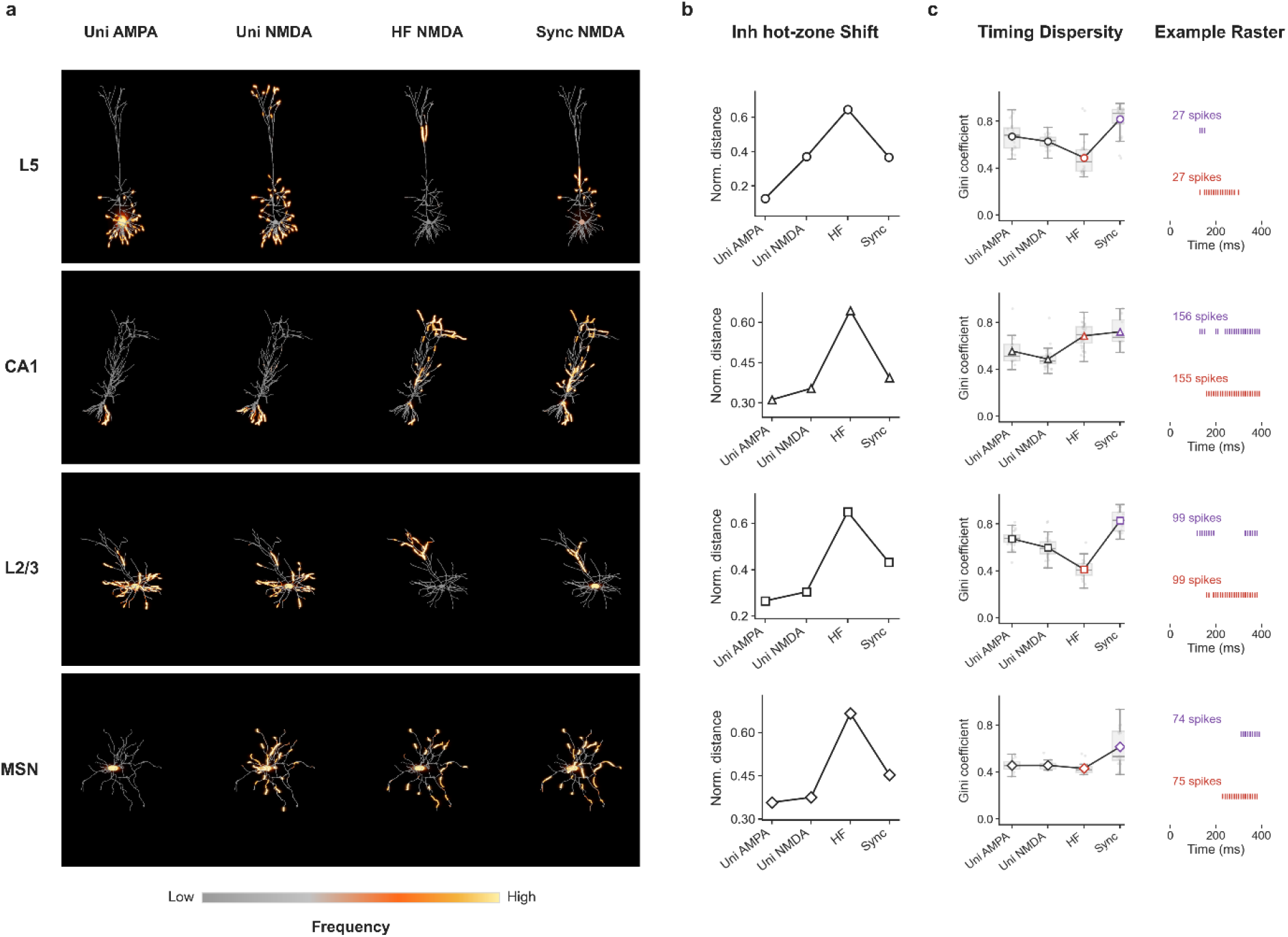
Dendritic inhibition follows a conserved spatial and temporal trajectory across neuron classes. (a) ODI maps for layer 5 (L5), CA1 and layer 2/3 (L2/3) pyramidal cells and striatal medium spiny neurons (MSNs) under uniform AMPA, uniform NMDA/AMPA, high-frequency distal NMDA/AMPA and synchronous distal NMDA/AMPA excitation^21–25^. Warm colours indicate higher selected-inhibition frequency. (b) Normalized distance of the inhibitory hot zone from the soma across excitation regimes. Across neuron classes, weak distributed excitation favours proximal or basal inhibition, high-frequency clustered excitation pushes inhibition distally, and synchronous clustered excitation retracts inhibition toward intermediate sites. (c) Temporal dispersion of selected inhibitory events quantified by the Gini coefficient. Synchronous clustered excitation produces more temporally clustered inhibition than high-frequency clustered excitation. (d) Representative inhibitory rasters showing selected inhibition timings for HF NMDA and Sync NMDA across models. Tick marks indicate individual selected inhibitory events.

The response-law decomposition revealed, however, that this shared macroscopic strategy was implemented through different biophysical mechanisms. In electrically compartmentalized neurons, exemplified by layer 5 and CA1 pyramidal cells, the susceptibility kernel **Φ**_*s*_ was strongly reshaped by excitation pattern. Distal regenerative events substantially altered the global susceptibility landscape, allowing remote or intermediate compartments to gain strong somatic leverage (Fig. 6b, bottom). In these cells, spatial remapping of inhibition was driven jointly by changes in global susceptibility and local synaptic drive (Fig. 6d, bottom).

**Fig. 6.**
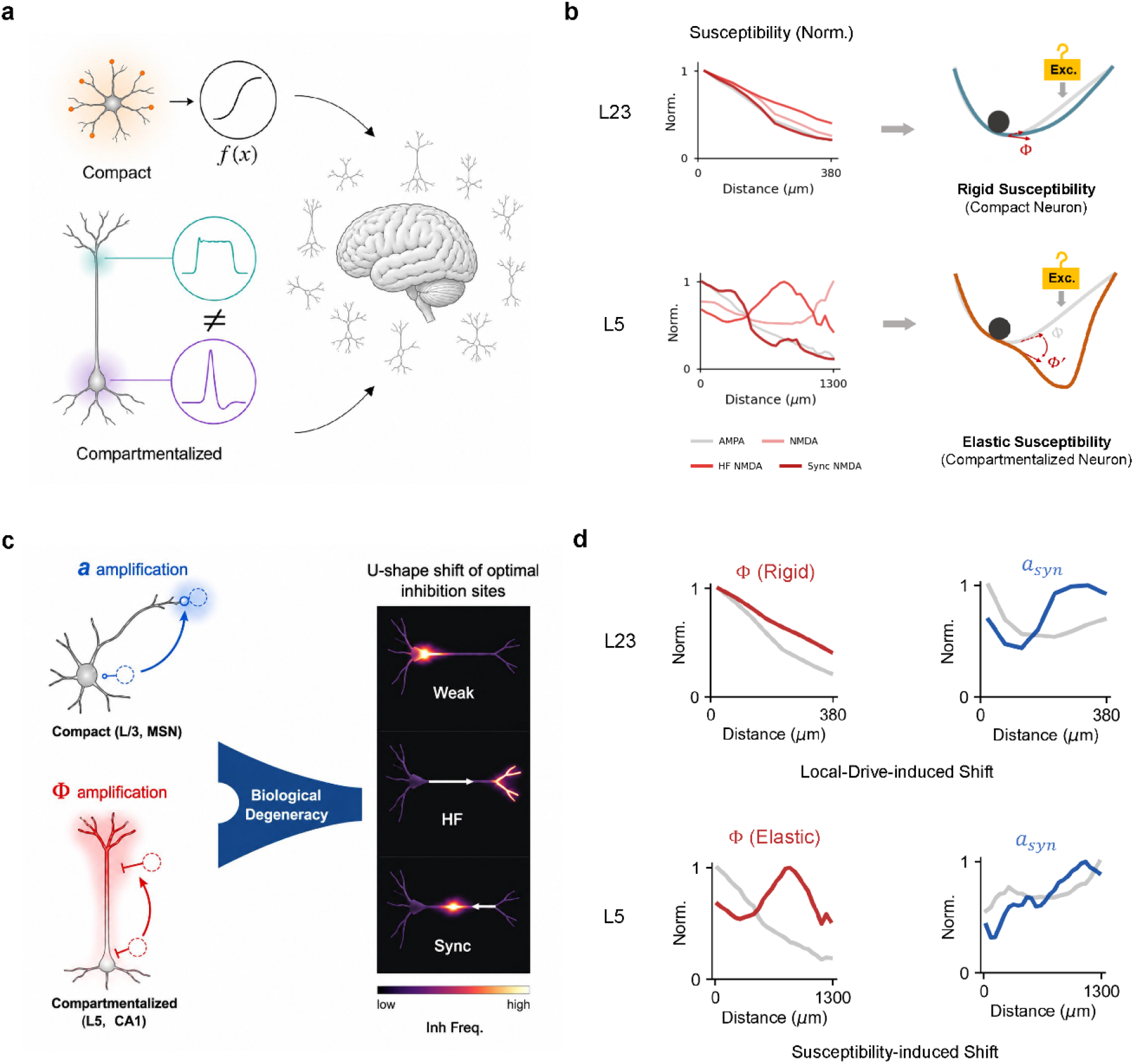
Compact and compartmentalized neurons implement the same inhibitory logic through different terms of the Φ–a law. (a) Conceptual contrast between compact and compartmentalized dendritic architectures^5,12,38^. Compact neurons can be described by a relatively rigid input–output nonlinearity, whereas compartmentalized neurons generate distinct dendritic events in different subcellular domains. (b) Normalized susceptibility profiles for representative compact (L2/3) and compartmentalized (L5) neurons across excitation regimes. Compact neurons retain a broadly distance-dependent, rigid susceptibility profile, whereas compartmentalized neurons show excitation-dependent susceptibility reshaping. (c) Mechanistic degeneracy of inhibitory remapping. Compact neurons shift inhibitory leverage mainly through local-drive amplification (*a*_*syn*_) at stimulated dendrites, whereas compartmentalized neurons shift leverage through amplification and reconfiguration of global susceptibility (Φ). Both mechanisms can generate a similar U-shaped shift in optimal inhibitory sites. (d) Response-law decomposition in representative L2/3 and L5 cells. In L2/3 cells, Φ changes little across excitation regimes and spatial remapping is dominated by *a*_*syn*_; in L5 cells, Φ is strongly reshaped by structured excitation, producing susceptibility-driven remapping of inhibitory hot zones.

In more electrically compact neurons, exemplified by layer 2/3 pyramidal cells and medium spiny neurons, the normalized susceptibility profile remained comparatively stable across excitation regimes and continued to decrease broadly with distance (Fig. 6b, top). In these cells, the shift in optimal inhibition was associated more strongly with state-dependent increases in the local perturbation term ***a***_*syn*_ at the stimulated compartments (Fig. 6d, top). Thus, compact neurons implement similar macroscopic inhibitory reorganization with less deformation of the global susceptibility landscape.

Morphologically and biophysically distinct neuron classes can therefore implement a shared inhibitory logic by exploiting different terms of the same response law. Compartmentalized neurons remap inhibition largely by reshaping global susceptibility, whereas compact neurons rely more strongly on local amplification of synaptic drive. This functional convergence through mechanistic divergence provides a quantitative example of biological degeneracy in dendritic computation (Fig. 6c).

## Discussion

The integration of excitatory and inhibitory (E/I) inputs within active dendrites is central to neural computation^5, 38^. Diverse GABAergic interneurons target specific subcellular domains of principal neurons, ranging from perisomatic compartments to distal dendrites, yet how this spatial diversity implements general rules of inhibitory control remains unclear^8-10^. The biophysical complexity of active dendrites has motivated detailed compartmental modeling, in which cable equations and membrane mechanisms are discretized across neuronal morphology and solved numerically^4, 31, 39^. Such models provide powerful in-silico tools for testing mechanistic hypotheses, but their high dimensionality, parameter richness and simulator-oriented formulation have limited their use for direct analytical theory^40-42^.

To reveal the general rule of dendritic inhibition, our work directly faced the intrinsic complexity of detailed neuron models and developed an analytical framework for investigating spatiotemporal strategies of effective inhibition in arbitrary neurons. We first reformulated the implicit simulation of detailed models into iterative matrix equations and derived the corresponding recursion of voltage gradients, which capture first-order perturbation dynamics with respect to synaptic weights. These gradients enabled us to identify efficient inhibitory strategies under arbitrary excitatory drives. Importantly, under structured high-frequency NMDA input onto distal dendrites, we observed a robust shift of inhibitory hot zones from perisomatic regions toward distal dendritic compartments. This predicted shift mirrors experimentally observed activity-dependent routing of inhibition in recurrent microcircuits, where sustained or burst-like pyramidal activity preferentially recruits dendrite-targeting inhibitory pathways rather than only perisomatic inhibition^43 44-47^. This consistency suggests that recurrent inhibitory motifs may be organized to implement efficient, state-dependent control of active dendritic excitation.

Moreover, we provided mechanistic explanations for emerging spatiotemporal inhibition patterns through the decomposition of voltage gradients: inhibitory effectiveness is jointly determined by the intrinsic global susceptibility of active dendrites and the local voltage response to synaptic perturbations. Interestingly, the spatiotemporal features of effective dendritic inhibition exhibit similar changes across all neurons examined when excitation patterns are altered, whereas the mechanisms underlying these changes diverge between compartmentalized and compact neurons, indicating both the universality and diversity of dendritic inhibition.

### Unification and expansion of classic theory on dendritic inhibition

Classical studies of dendritic inhibition found the shunting effect reaches its maximum when inhibition is located on the path from the excitation to the soma^26^, while distal inhibition surpasses on-path inhibition when clustered NMDA inputs are presented^13^. Since 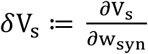 can well approximate the actual somatic impact ΔV_s_ especially for smooth voltage responses from passive models, the prediction of our algorithm when only considering the first inhibitory input should perfectly match previous results mentioned above. Nevertheless, our predictions seem contradictory to previous findings: the most efficient inhibition sites for pure uniform AMPA contain basal terminals apart from perisomatic compartments. The main reason is that we used active neuron models where the membrane conductance of the soma is significantly larger than passive models’, and on-site/off-path inhibition is stronger than proximal inhibition regardless of NMDA inputs if somatic membrane conductance is sufficiently large (Methods). The second reason is ODI requires finding a group of inhibitory inputs to suppress somatic spikes, and the aggregation of inhibitory inputs at one dendritic domain will lead to the saturation of inhibition, thus preventing pure perisomatic inhibition. Interestingly, our prediction shows that the portion of perisomatic inhibition will increase when considering fast-spiking GABA inputs near the soma (path distance < 50 *μm*) for unstructured excitations, while remaining unchanged for distal structured excitations, indicating different roles played by perisomatic and distal inhibition (Supplementary Fig. 4).

### Global susceptibility: the novel metric quantifying the kinetic divergence across different excitation patterns and neuron types

One of the long-pursued goals in theoretical neuroscience is to compress computational properties of realistic neurons into one general formulation. Since transfer impedances between dendrites and the soma fluctuate greatly with membrane voltages on active neurons^14-16^, static descriptions of this I/O process are fundamentally untenable. In this study, we provided a dynamic expression of dendritic computation through formulating the neuronal response to synaptic perturbations, instead of sticking on classical I/O functions. We proved the first-order somatic output, *∂V*_*s*_, is the inner product of two vectors corresponding to two distinct aspects of dendritic properties. Specifically, the intrinsic susceptibility Φ_s_ reflects how synaptic perturbations spread globally to influence the somatic response, independent of the perturbation amplitude. Therefore, Φ_s_ is the dynamic snapshot of the neuron I/O function at the first-order domain, which exhibits different levels of elasticity by external excitations among different neuron types. Interestingly, for neurons with longer electrical distance and more elaborate arborizations of distal dendrites, e.g. L5 PC and CA1 PC, Φ_s_ at distal domains can be significantly elevated by strong distal excitations, while Φ_s_ remains monotonically decreasing from soma to distal domains for neurons with more compact electrical structures (L2/3 PC, MSN). One intuitive explanation for this divergence is that for compartmentalized neurons, dendritic events such as Ca^2+^ plateau locally integrate and amplify the influence of distal inputs, with little impact on proximal inputs, and this highly-local elevation results in the excitation-dependent Φ_s_. For compact neurons, however, these dendritic events spread globally and equally influence input integration on the whole dendritic tree, leading to synchronous up/down regulations of Φ_s_ at distinct domains. Further analyses on the quantitative relationship between Φ_s_ and electrical parameters of neurons are required to consolidate this postulation. Nonetheless, the global susceptibility Φ_s_ successfully quantifies the kinetic diversity among different neuron types and potentially guides categorizations according to the electrical properties of neurons.

### Across-scale linkage between cognitive functions and dendritic inhibition

The balance of excitatory glutamatergic inputs and inhibitory GABAergic inputs is crucial for signal integration and information processing in neuronal microcircuits, and disruption of E/I balance at cellular and network levels is strongly associated with cognitive and neuropsychiatric dysfunctions^48-50^. Specifically, somatostatin-positive interneurons (SST-INs) preferentially target distal dendritic compartments of principal neurons and regulate the gating and integration of dendritic inputs through local and recurrent circuit recruitment^10, 44, 51^. Consistent with this circuit role, reduced SST signaling has been repeatedly implicated in neuropsychiatric and neurodegenerative disorders^52, 53^. For example, SST expression is reduced in the dorsolateral PFC^54^, subgenual/anterior cingulate cortex^55, 56^ and amygdala^57^ of major depressive disorder (MDD), whereas PV-related changes appear less consistent in MDD^54^. Related SST-interneuron dysfunction has also been reported in Alzheimer’s disease (AD). In hippocampal circuits, oriens lacunosum-moleculare (O-LM) interneurons regulate CA1 input routing, and AD-like beta-amyloidosis impairs cholinergic drive and synaptic rewiring of O-LM interneurons, linking SST-positive interneuron dysfunction to memory deficits^58-60^. While these experimental findings support the importance of apical dendritic inhibition in feedback-related cognitive functions^61^, the underlying mechanisms at the level of dendritic computation remain difficult to isolate experimentally in behaving animals. Our theoretical approach predicted the spatiotemporal features of efficient dendritic inhibition for structured excitations on distal apical dendrites that mimic feedback inputs, aiding the design of experiments to link dendritic inhibition to cognitive functions, for example by blocking GABAergic inputs at predicted sites for optimal inhibition and examining the influence of the blockade on animal behavior. Moreover, our approach can be further generalized to investigate E/I integration in neuronal microcircuits by expanding our algorithm to enable network-level gradient computations, serving as our next step toward a unified, multiscale mechanism of E/I balance in the neuronal system.

## Methods

### Spatial Inhibitory Efficacy on Passive Dendrites

To systematically investigate the spatial determinants of inhibitory synaptic efficacy, we started from the steady-state analytical approach using a classical “ball-and-stick” neuronal model. The model consists of a somatic compartment with a lumped leak conductance *G*_*s*_, coupled to a uniform, passive cylindrical dendrite of electrotonic length *L* and semi-infinite input impedance *G*_∞_. An excitatory synapse (conductance *g*, reversal potential *E*) was positioned at an electrotonic distance *l* from the soma, while an inhibitory synapse (conductance *g*_*i*_, reversal potential *E*_*i*_ = 0) was introduced at a variable location *l*_*i*_. Inhibitory efficacy was quantified by the degree to which it minimized the steady-state somatic voltage *V*(0, *l*_*i*_).

We analyzed the relative suppressive strengths of somatic (*l*_*i*_ = 0), on-path/proximal (0 < *l*_*i*_ < *l*), on-site (*l*_*i*_ = *l*), and off-path/distal (*l*_*i*_ > *l*) inhibition. Classical theoretical studies often assert that proximal, on-path inhibition is universally the most effective at shunting somatic voltage. However, our exact analytical solutions reveal that this hierarchy is strictly dependent on somatic and dendritic parameters. Specifically, for proximal inputs (*l*_*i*_ ≤ *l*), the superiority of somatic or on-path inhibition implicitly assumes a low somatic resting conductance. We establish that somatic inhibition is only strictly stronger than on-site inhibition when the following inequality holds:

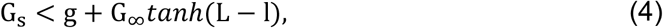

If *G*_*s*_ exceeds this critical threshold—a physiologically relevant scenario during the opening of voltage-gated sodium channels near the spike threshold—the classical hierarchy breaks down, and on-site or even off-path inhibition can theoretically outperform proximal on-path inhibition. Conversely, for distal off-path inhibition (*l*_*i*_ > *l*), spatial efficacy exhibits a simpler profile. The absolute magnitude of the somatic voltage gradient with respect to the inhibitory conductance is a monotonically decreasing function of the electrotonic distance from the soma. Thus, for relatively small inhibitory conductances, the efficacy of off-path inhibition strictly decays as the physical distance from the excitatory site increases.

The revelation that optimal inhibitory placement is highly sensitive to localized parameters like somatic conductance and excitation strength within a simplified continuous model highlights a critical theoretical bottleneck. As analyses scale to encompass realistic neuronal complexities—such as intricate dendritic branching geometries, transient synaptic conductances, and active voltage-gated ion channels—deriving exact analytical solutions via continuous partial differential equations becomes mathematically intractable. This fundamental limitation of continuous cable theory necessitates a paradigm shift toward a robust and scalable numerical framework. Consequently, this motivated our development of a generalizable matrix formulation for detailed multi-compartment models, allowing for the precise computation of E/I interactions across arbitrary morphologies (Detailed continuous derivations are provided in Supplementary Note 1).

### Derivation of perturbation kinetics and the response law on realistic neuron models

Membrane voltages were updated using the standard implicit discretization implemented in NEURON^32^. Starting from the discretized cable equation with voltage-dependent membrane mechanisms, one simulation step can be written as a linear system for the next-step membrane voltage. After eliminating auxiliary algebraic variables introduced by the implicit discretization, this system can be reduced to an affine recursion of the form

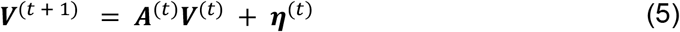

where **V**^(t)^ is the membrane-voltage vector over all physical compartments. Here, **A**^(t)^ and **η**^(t)^ are both evaluated on the current trajectory and therefore depend implicitly on the current voltage state.

For a candidate inhibitory input *syn*, we then considered the first-order response of the voltage trajectory to its synaptic weight. Because both **A**^(t)^ and **η**^(t)^ depend on the current voltage state, the perturbation dynamics is governed not by **A**^(t)^ itself, but by the Jacobian of the one-step voltage update map. Denoting the voltage response to unit perturbation on the synaptic weight *w*_*syn*_ as 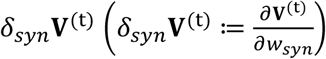 in recursion can be written as

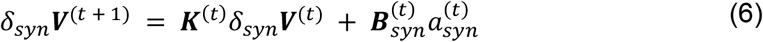

where **K**^(t)^ is the Jacobian propagator of the one-step voltage map evaluated on the current trajectory, 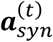 is the effective local perturbation waveform of the candidate inhibitory input, and 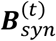 specifies how a unit perturbation of *syn* enters the full voltage system after the implicit update has been resolved.

Assuming zero initial perturbation, repeated substitution yields the voltage response at readout time t*:

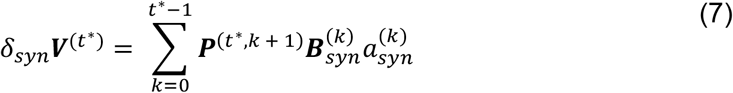

where

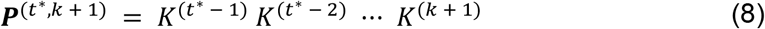

is the transition operator generated by the successive Jacobian propagators.

Projecting this response onto the soma using the soma readout vector ***e***_*s*_ gives

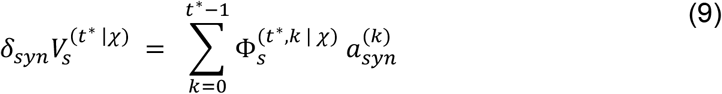

with

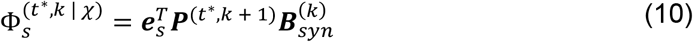

Thus, for a fixed candidate inhibitory input and a fixed current state, the somatic effect is decomposed into the inner product between a local perturbation waveform and a trajectory-dependent sensitivity vector. In the compact notation used in the main text,

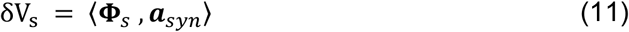

the readout time *t*^∗^ and current trajectory *χ* are suppressed for clarity, and the relevant temporal degrees of freedom are written in vectorized form.

Because the dendritic system is nonlinear, **Φ**_s_ depends on the current background trajectory and is recomputed after each accepted inhibitory perturbation. Therefore, this compact formula describes a local first-order effect around the current state, rather than a global linear superposition across multiple inhibitory inputs. Full algebraic details are provided in Supplementary Note 1.

### Formal definition and complexity proof for optimal dendritic inhibition

We considered a deterministic multi-compartment neuron model simulated over a finite discrete time horizon. The dendritic tree was discretized into a finite set of candidate inhibitory locations, and time was discretized into simulation steps. For a fixed excitatory input pattern, let *P* denote the set of all candidate inhibitory pulses across space and time. For any subset *S* ⊆ *P*, the somatic voltage trace generated by the simulator is denoted by *V*(*t*; *S*). We say that *S* is successful if the neuron does not emit a somatic spike during the observation window, that is, if the somatic voltage remains below threshold for all time points.

The optimization problem is to identify the smallest inhibitory subset *S* that prevents somatic spiking. Its associated decision form asks whether such a subset exists with size at most *K*. This decision problem belongs to NP, because once a candidate subset *S* is given, its validity can be checked by one forward simulation followed by threshold detection on the somatic voltage trace.

To establish hardness, we considered a monotone-inhibition subclass in which adding extra inhibitory pulses cannot increase the somatic voltage at any time point. This assumption captures the intuitive regime in which stronger or additional inhibition cannot paradoxically make the neuron more excitable. We then reduced the classical Set Cover problem to ODI. Given a Set Cover instance with universe *U* = {*u*_1_, …, *u*_*s*_}, family of sets 𝒞 = {*C*_1_, …, *C*_*m*_}, and budget *K*, we constructed a neuron-model instance as follows. The full simulation window was partitioned into *t* non-overlapping temporal windows, one for each element *u*_*i*_ ∈ *U*. In the absence of inhibition, each window produced an isolated suprathreshold somatic event. These windows were separated by sufficient recovery intervals so that they behaved independently. We then introduced *μ* candidate inhibitory pulses, one for each set *C*_*j*_. The pulse corresponding to *C*_*j*_ was defined so that it suppressed the spike in window *W*(*u*_*i*_) if and only if *u*_*i*_ ∈ *C*_*j*_, while leaving the other windows unsuppressed.

Under monotone inhibition, a collection of inhibitory pulses suppresses all spike windows exactly when the corresponding collection of sets covers all elements of the universe. Therefore, the Set Cover instance has a solution of size at most *K* if and only if the constructed ODI instance has a spike-suppressing inhibitory subset of size at most *K*. Because the construction requires only a polynomial number of temporal windows, candidate pulses, and model parameters, this gives a polynomial-time reduction from Set Cover to ODI.

It follows that the decision version of optimal dendritic inhibition is NP-complete, and that the corresponding minimization problem is NP-hard. In other words, the difficulty of finding the globally minimal inhibitory pattern is intrinsic to the combinatorial structure of the problem, rather than an artifact of brute-force search. This complexity result motivates the use of approximate procedures in the main text, including the gradient-based strategy developed here.

### Algorithm validation on the toy neuron model

To validate the algorithm in a controlled setting, we used the classical ball-and-stick neuron model implemented in NEURON, consisting of a one-compartment soma and a three-compartment dendrite. The soma had length and diameter *L*_*soma*_ = *d*_*soma*_ = 20 *μ*m, and the dendrite had length *L*_*dend*_ = 60 *μ*m and diameter *d*_*dend*_ = 1 *μ*m. All compartments contained Hodgkin–Huxley channels with default parameters (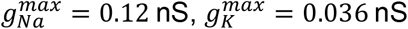, and *g*_*l*_ = 0.00003 nS).

The total simulation duration was 60 ms. No external inputs were applied during the first 20 ms, allowing the membrane potential to reach a steady state before stimulus onset. Excitatory drive was provided by NMDA synapses placed at random dendritic locations, each with fixed conductance *g*_*NMDA*_ = 0.001 nS, rise time constant 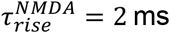, and decay time constant 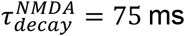. Each NMDA synapse fired once, with spike time uniformly sampled from the interval [10 ms, 60 ms].

At each step of the algorithm, inhibition was introduced as a GABA pulse with fixed conductance *g*_*GABA*_ = 0.001 nS, rise time constant 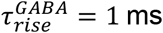, and decay time constant 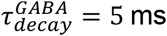. The candidate spike time of each GABA pulse was restricted to the discrete set [20, 30, 40, 50, 60] ms.

For each test case, the optimal inhibitory solution was determined by brute-force search. To control the computational cost of exhaustive enumeration, we searched only subsets containing no more than 10 inhibitory spikes and retained cases for which the ground-truth optimal solution could be identified within this search range. Algorithm-derived solutions were then compared with the corresponding ground-truth solutions. The comparison was visualized as a two-dimensional heatmap (Fig. 2f), in which each grid cell represents the frequency of a given algorithm-solution/ground-truth pair.

### Excitation pattern design

Each simulation consisted of a 100 ms background-input period followed by 300 ms of pure excitatory drive. Across all four excitation patterns, both excitatory and inhibitory synapses were modeled as two-exponential conductance-based synapses driven by Poisson spike trains. Background activity consisted of AMPA inputs firing at approximately 1 Hz and GABA inputs firing at approximately 10 Hz. The maximal conductances were set to 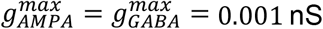. The AMPA time constants were 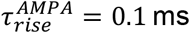 and 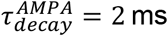, and the GABA time constants were 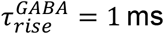 and 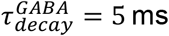. The ratio *N*_*AMPA*_: *N*_*GABA*_ was set to 4:1 to mimic an E/I-balanced regime in realistic neural circuits, while the absolute numbers of AMPA and GABA synapses were adjusted across excitation patterns and neuron types to maintain a moderate excitation level. Only excitatory inputs were presented during the interval from 100 ms to 400 ms.

For the **Uniform AMPA** pattern, all excitatory inputs were AMPA synapses with the same conductance and time constants as the background AMPA inputs, and only the total number of inputs was adjusted to generate somatic spikes.

For the **Uniform NMDA/AMPA** pattern, NMDA and AMPA synapses were combined at a default ratio of 1:1. NMDA synapses had 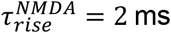 and 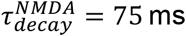. Uniformly distributed excitatory inputs in this condition fired at approximately 3 Hz.

For the **High-Frequency Distal Cluster** pattern, excitatory input consisted of two components: uniformly distributed NMDA/AMPA inputs firing at approximately 1 Hz, and clustered NMDA/AMPA inputs located on distal dendrites firing at approximately 10 Hz. Synapses belonging to the same cluster were positioned within 20 *μ*m of the cluster center.

For the **Synchronous Distal Cluster** pattern, excitatory input again consisted of uniformly distributed NMDA/AMPA inputs together with a distal clustered component. The uniformly distributed NMDA/AMPA inputs fired at approximately 5 Hz. Each synchronous cluster consisted of 20 NMDA/AMPA synapses activated sequentially within a short 2 ms time window at the same dendritic location, resulting in an extremely high instantaneous firing rate.

## Code availability

The code used for simulations and analyses will be made available upon reasonable request.

## Funding

This work is supported by National Natural Science Foundation of China grants (32471149).

## Competing interests

The authors declare no competing interests.

## Notes

### Competing Interest Statement

The authors have declared no competing interest.

## References

1. Isaacson, J.S. & Scanziani, M. How inhibition shapes cortical activity. Neuron 72, 231–243 (2011).

2. Vogels, T.P., Sprekeler, H., Zenke, F., Clopath, C. & Gerstner, W. Inhibitory plasticity balances excitation and inhibition in sensory pathways and memory networks. Science 334, 1569–1573 (2011).

3. Xue, M., Atallah, B.V. & Scanziani, M. Equalizing excitation-inhibition ratios across visual cortical neurons. Nature 511, 596–600 (2014).

4. Rall, W. Theory of physiological properties of dendrites. Annals of the New York Academy of Sciences 96, 1071–1092 (1962).

5. London, M. & Hausser, M. Dendritic computation. Annual Review of Neuroscience 28, 503–532 (2005).

6. Stuart, G.J. & Spruston, N. Dendritic integration: 60 years of progress. Nature Neuroscience 18, 1713–1721 (2015).

7. Silver, R.A. Neuronal arithmetic. Nature Reviews Neuroscience 11, 474–489 (2010).

8. Markram, H., et al. Interneurons of the neocortical inhibitory system. Nature Reviews Neuroscience 5, 793–807 (2004).

9. Kepecs, A. & Fishell, G. Interneuron cell types are fit to function. Nature 505, 318–326 (2014).

10. Tremblay, R., Lee, S. & Rudy, B. GABAergic interneurons in the neocortex: from cellular properties to circuits. Neuron 91, 260–292 (2016).

11. Schiller, J., Major, G., Koester, H.J. & Schiller, Y. NMDA spikes in basal dendrites of cortical pyramidal neurons. Nature 404, 285–289 (2000).

12. Major, G., Larkum, M.E. & Schiller, J. Active properties of neocortical pyramidal neuron dendrites. Annual Review of Neuroscience 36, 1–24 (2013).

13. Gidon, A. & Segev, I. Principles governing the operation of synaptic inhibition in dendrites. Neuron 75, 330–341 (2012).

14. Destexhe, A., Rudolph, M. & Pare, D. The high-conductance state of neocortical neurons in vivo. Nature Reviews Neuroscience 4, 739–751 (2003).

15. Ulrich, D. Dendritic resonance in rat neocortical pyramidal cells. Journal of Neurophysiology 87, 2753–2759 (2002).

16. Yaron-Jakoubovitch, A., Jacobson, G.A., Koch, C., Segev, I. & Yarom, Y. A paradoxical isopotentiality: a spatially uniform noise spectrum in neocortical pyramidal cells. Frontiers in Cellular Neuroscience 2, 3 (2008).

17. Holt, G.R. & Koch, C. Shunting inhibition does not have a divisive effect on firing rates. Neural Computation 9, 1001–1013 (1997).

18. Chance, F.S., Abbott, L.F. & Reyes, A.D. Gain modulation from background synaptic input. Neuron 35, 773–782 (2002).

19. Prescott, S.A. & de Koninck, Y. Gain control of firing rate by shunting inhibition: Roles of synaptic noise and dendritic saturation. Proceedings of the National Academy of Sciences of the United States of America 100, 2076–2081 (2003).

20. Mitchell, S.J. & Silver, R.A. Shunting inhibition modulates neuronal gain during synaptic excitation. Neuron 38, 433–445 (2003).

21. Iascone, D.M., et al. Whole-Neuron Synaptic Mapping Reveals Spatially Precise Excitatory/Inhibitory Balance Limiting Dendritic and Somatic Spiking. Neuron 106, 566–578.e568 (2020).

22. Hay, E., Hill, S., Schurmann, F., Markram, H. & Segev, I. Models of neocortical layer 5b pyramidal cells capturing a wide range of dendritic and perisomatic active properties. PLoS Computational Biology 7, e1002107 (2011).

23. Poirazi, P., Brannon, T. & Mel, B.W. Arithmetic of subthreshold synaptic summation in a model CA1 pyramidal cell. Neuron 37, 977–987 (2003).

24. Poirazi, P., Brannon, T. & Mel, B.W. Pyramidal neuron as two-layer neural network. Neuron 37, 989–999 (2003).

25. Lindroos, R., et al. Basal Ganglia Neuromodulation Over Multiple Temporal and Structural Scales-Simulations of Direct Pathway MSNs Investigate the Fast Onset of Dopaminergic Effects and Predict the Role of Kv4.2. Frontiers in Neural Circuits 12, 3 (2018).

26. Koch, C., Poggio, T. & Torre, V. Nonlinear interactions in a dendritic tree: localization, timing, and role in information processing. Proceedings of the National Academy of Sciences of the United States of America 80, 2799–2802 (1983).

27. Vu, E.T. & Krasne, F.B. Evidence for a computational distinction between proximal and distal neuronal inhibition. Science 255, 1710–1712 (1992).

28. Hao, J., Wang, X.-d., Dan, Y., Poo, M.-m. & Zhang, X.-h. An arithmetic rule for spatial summation of excitatory and inhibitory inputs in pyramidal neurons. Proceedings of the National Academy of Sciences of the United States of America 106, 21906–21911 (2009).

29. Jadi, M., Polsky, A., Schiller, J. & Mel, B.W. Location-dependent effects of inhibition on local spiking in pyramidal neuron dendrites. PLoS Computational Biology 8, e1002550 (2012).

30. Pouille, F., Watkinson, O., Scanziani, M. & Trevelyan, A.J. The contribution of synaptic location to inhibitory gain control in pyramidal cells. Physiological Reports 1, e00067 (2013).

31. Hines, M.L. & Carnevale, N.T. The NEURON simulation environment. Neural Computation 9, 1179–1209 (1997).

32. Carnevale, N.T. & Hines, M.L. The NEURON Book (Cambridge University Press, 2006).

33. Waters, J. & Helmchen, F. Background Synaptic Activity Is Sparse in Neocortex. Journal of Neuroscience 26, 8267–8277 (2006).

34. Gasparini, S. & Magee, J.C. State-Dependent Dendritic Computation in Hippocampal CA1 Pyramidal Neurons. Journal of Neuroscience 26, 2088–2100 (2006).

35. Polsky, A., Mel, B.W. & Schiller, J. Computational Subunits in Thin Dendrites of Pyramidal Cells. Nature Neuroscience 7, 621–627 (2004).

36. Larkum, M.E., Zhu, J.J. & Sakmann, B. A new cellular mechanism for coupling inputs arriving at different cortical layers. Nature 398, 338–341 (1999).

37. Palmer, L.M., et al. NMDA Spikes Enhance Action Potential Generation during Sensory Input. Nature Neuroscience 17, 383–390 (2014).

38. Magee, J.C. Dendritic Integration of Excitatory Synaptic Input. Nature Reviews Neuroscience 1, 181–190 (2000).

39. Rall, W. Branching dendritic trees and motoneuron membrane resistivity. Experimental Neurology 1, 491–527 (1959).

40. Achard, P. & De Schutter, E. Complex parameter landscape for a complex neuron model. PLoS Computational Biology 2, e94 (2006).

41. Marder, E. & Taylor, A.L. Multiple models to capture the variability in biological neurons and networks. Nature Neuroscience 14, 133–138 (2011).

42. Almog, M. & Korngreen, A. Is realistic neuronal modeling realistic? Journal of Neurophysiology 116, 2180–2209 (2016).

43. Pouille, F.d.r. & Scanziani, M. Routing of spike series by dynamic circuits in the hippocampus. Nature 429, 717–723 (2004).

44. Silberberg, G. & Markram, H. Disynaptic inhibition between neocortical pyramidal cells mediated by Martinotti cells. Neuron 53, 735–746 (2007).

45. Berger, T.K., Perin, R., Silberberg, G. & Markram, H. Frequency-dependent disynaptic inhibition in the pyramidal network: a ubiquitous pathway in the developing rat neocortex. The Journal of Physiology 587, 5411–5425 (2009).

46. Berger, T.K., Silberberg, G., Perin, R. & Markram, H. Brief bursts self-inhibit and correlate the pyramidal network. PLoS Biology 8, e1000473 (2010).

47. Murayama, M., et al. Dendritic encoding of sensory stimuli controlled by deep cortical interneurons. Nature 457, 1137–1141 (2009).

48. Yizhar, O., et al. Neocortical excitation/inhibition balance in information processing and social dysfunction. Nature 477, 171–178 (2011).

49. Denève, S. & Machens, C.K. Efficient codes and balanced networks. Nature Neuroscience 19, 375–382 (2016).

50. Sohal, V.S. & Rubenstein, J.L.R. Excitation-inhibition balance as a framework for investigating mechanisms in neuropsychiatric disorders. Molecular Psychiatry 24, 1248–1257 (2019).

51. Urban-Ciecko, J. & Barth, A.L. Somatostatin-expressing neurons in cortical networks. Nature Reviews Neuroscience 17, 401–409 (2016).

52. Lin, L.-C. & Sibille, E. Reduced brain somatostatin in mood disorders: a common pathophysiological substrate and drug target? Frontiers in Pharmacology 4, 110 (2013).

53. Fee, C., Banasr, M. & Sibille, E. Somatostatin-Positive Gamma-Aminobutyric Acid Interneuron Deficits in Depression: Cortical Microcircuit and Therapeutic Perspectives. Biological Psychiatry 82, 549–559 (2017).

54. Sibille, E., Morris, H.M., Kota, R.S. & Lewis, D.A. GABA-related transcripts in the dorsolateral prefrontal cortex in mood disorders. The International Journal of Neuropsychopharmacology 14, 721–734 (2011).

55. Tripp, A., Kota, R.S., Lewis, D.A. & Sibille, E. Reduced somatostatin in subgenual anterior cingulate cortex in major depression. Neurobiology of Disease 42, 116–124 (2011).

56. Seney, M.L., Tripp, A., McCune, S., Lewis, D.A. & Sibille, E. Laminar and cellular analyses of reduced somatostatin gene expression in the subgenual anterior cingulate cortex in major depression. Neurobiology of Disease 73, 213–219 (2015).

57. Guilloux, J.-P., et al. Molecular evidence for BDNF- and GABA-related dysfunctions in the amygdala of female subjects with major depression. Molecular Psychiatry 17, 1130–1142 (2012).

58. Leão, R.N., et al. OLM interneurons differentially modulate CA3 and entorhinal inputs to hippocampal CA1 neurons. Nature Neuroscience 15, 1524–1530 (2012).

59. Palop, J.J. & Mucke, L. Network abnormalities and interneuron dysfunction in Alzheimer disease. Nature Reviews Neuroscience 17, 777–792 (2016).

60. Schmid, L.C., et al. Dysfunction of Somatostatin-Positive Interneurons Associated with Memory Deficits in an Alzheimer’s Disease Model. Neuron 92, 114–125 (2016).

61. Manita, S., et al. A Top-Down Cortical Circuit for Accurate Sensory Perception. Neuron 86, 1304–1316 (2015).

