## Supplementary Methods and Figures for "A unified law for inhibitory control in active dendrites"

### 1. Passive-cable derivation for the spatial efficacy of a single inhibitory input

#### Model definition

We considered a steady-state passive ball-and-stick model in electrotonic coordinates x ∈ [0, L], with the soma located at x = 0 and a sealed dendritic end at x = L.[1] A single excitatory conductance g, with reversal potential E, was placed at x = l, and a single inhibitory conductance $g_i$, with reversal potential $E_i = 0$, was placed at $x = l_i$. The resting potential was set to 0. The input conductance of a semi-infinite passive cable with the same passive parameters is denoted by $G_\infty$.

Between synaptic discontinuities, the steady-state membrane potential satisfies the passive cable equation

$$\frac{d^2V}{dx^2} = V. \tag{S1}$$

Therefore, on each cable segment, the general solution takes the form

$$V(x) = A\sinh x + B\cosh x. \tag{S2}$$

The coefficients are determined by imposing voltage continuity and axial-current conservation at the synaptic positions, together with the sealed-end condition at x = L and the lumped somatic conductance at x = 0.

#### Closed-form expression for the somatic voltage

For inhibition located between the soma and the excitatory input ($l_i \leq l$), the somatic voltage is

$$V_s \equiv V(0; l_i) = \frac{gE}{g_i F_1(l_i) + C_1(l, L)}. \tag{S3}$$

Here

$$\begin{aligned} F_1(l_i) = {} & \frac{G_s}{G_\infty^2}(g + G_\infty)\sinh l_i \sinh(l - l_i) \\ & + \cosh l_i \cosh(l - l_i) \\ & + \left(\frac{g}{G_\infty} + \tanh(L - l)\right)\cosh l_i \sinh(l - l_i) \end{aligned} \tag{S4}$$

$$+\frac{G_s}{G_\infty}\sinh l_i \cosh(l - l_i).$$

and

$$C_1(l, L) = \big(G_s + g + G_\infty \tanh(L - l)\big)\cosh l + \left(G_\infty + \frac{g G_s}{G_\infty} + G_s \tanh(L - l)\right)\sinh l. \tag{S5}$$

For inhibition located distal to the excitatory input ($l_i > l$), the somatic voltage becomes

$$V_s \equiv V(0; l_i) = \frac{\big(G_\infty + g_i F_2(l_i)\big) g E}{g_i F_3(l_i) + C_2(l, L)}. \tag{S6}$$

The auxiliary functions are

$$F_2(l_i) = \frac{\tanh(l_i - l)}{1 + \tanh(L - l_i)\tanh(l_i - l)}. \tag{S7}$$

$$F_3(l_i) = \frac{f_1(l)\tanh(l_i - l) + f_2(l)}{1 + \tanh(L - l_i)\tanh(l_i - l)}. \tag{S8}$$

$$C_2(l, L) = G_\infty\big(f_1(l) + f_2(l)\tanh(L - l)\big). \tag{S9}$$

with

$$f_1(l) = \left(1 + \frac{g G_s}{G_\infty^2}\right)\sinh l + \frac{G_s + g}{G_\infty}\cosh l. \tag{S10}$$

$$f_2(l) = \frac{G_s}{G_\infty}\sinh l + \cosh l. \tag{S11}$$

Eqs. (S3)-(S11) are algebraically equivalent to the original derivation, but are rewritten here using a unified notation and explicit grouping of terms.

**Position dependence of inhibitory efficacy**

We define three geometric regimes relative to the excitatory site at $x = l$: soma-proximal on-path inhibition ($0 \le l_i < l$), on-site inhibition ($l_i = l$), and distal off-path inhibition ($l_i > l$).

For $l_i \le l$, minimizing the somatic voltage in Eq. (S3) is equivalent to maximizing $F_1(l_i)$. The two boundary values are

$$F_1(0) = \cosh l + \left(\frac{g}{G_\infty} + \tanh(L - l)\right)\sinh l. \tag{S12}$$

and

$$F_1(l) = \cosh l + \frac{G_s}{G_\infty}\sinh l. \tag{S13}$$

Therefore,

$$F_1(0) > F_1(l) \Leftrightarrow G_s < g + G_\infty \tanh(L - l). \tag{S14}$$

whereas

$$F_1(l) > F_1(0) \Leftrightarrow G_s > g + G_\infty \tanh(L - l). \tag{S15}$$

Eq. (S14) shows that the classical preference for proximal on-path inhibition is not universal, but depends on a specific relation among somatic conductance, excitatory strength, and dendritic length.[2,3] Once the effective somatic conductance exceeds the threshold in Eq. (S15), the strongest inhibitory position shifts to the excitatory site itself.

For $l_i > l$, the exact expression in Eq. (S6) is more cumbersome, but the weak-inhibition ranking is directly obtained from the voltage gradient evaluated at $g_i = 0$:

$$\left.\frac{\partial V_s}{\partial g_i}\right|_{g_i=0} = gE\,\frac{C_2(l,L)F_2(l_i) - G_\infty F_3(l_i)}{C_2(l,L)^2}. \tag{S16}$$

The prefactor gE is positive and therefore does not affect the spatial ranking. In this regime, $|\partial V_s/\partial g_i|$_{$g_i$=0} decreases monotonically with increasing $l_i$, indicating that the strongest off-path inhibition is attained in the limit $l_i \to l^+$, that is, immediately distal to the excitatory site. Accordingly, when $G_s$ is sufficiently large, both on-site inhibition and sufficiently near-site off-path inhibition can exceed the efficacy of classical proximal on-path inhibition.

### First-order approximation of finite voltage suppression

The weak-inhibition analysis above ranks inhibitory locations using the voltage gradient at $g_i = 0$. To quantify how accurately this gradient predicts the actual voltage reduction at finite inhibitory strength, we compared the exact voltage change,

$$\Delta V_{\text{exact}}(l_i; g_i) = V_s(l_i; g_i) - V_s(l_i; 0), \tag{S17}$$

with its first-order approximation,

$$\Delta V_{\text{lin}}(l_i; g_i) = \left.\frac{\partial V_s(l_i; g_i)}{\partial g_i}\right|_{g_i=0} g_i. \tag{S18}$$

For $l_i \le l$, Eq. (S3) gives

$$\Delta V_{\text{exact}}^{\text{prox}} = -\frac{gE\, g_i F_1(l_i)}{C_1(l,L)\big(C_1(l,L) + g_i F_1(l_i)\big)}. \tag{S19}$$

whereas the linear approximation is

$$\Delta V_{\text{lin}}^{\text{prox}} = -\frac{gE\, g_i F_1(l_i)}{C_1(l,L)^2}. \tag{S20}$$

Thus,

$$\Delta V_{\text{exact}}^{\text{prox}} = \Delta V_{\text{lin}}^{\text{prox}} \left(1 + \frac{g_i F_1(l_i)}{C_1(l,L)}\right)^{-1}. \tag{S21}$$

For $l_i > l$, Eq. (S6) yields

$$\Delta V_{\text{exact}}^{\text{dist}} = gE\, g_i \, \frac{C_2(l,L)F_2(l_i) - G_\infty F_3(l_i)}{C_2(l,L)\big(C_2(l,L) + g_i F_3(l_i)\big)}. \tag{S22}$$

and the corresponding first-order approximation is

$$\Delta V_{\text{lin}}^{\text{dist}} = gE\, g_i \, \frac{C_2(l,L)F_2(l_i) - G_\infty F_3(l_i)}{C_2(l,L)^2}. \tag{S23}$$

Therefore,

$$\Delta V_{\text{exact}}^{\text{dist}} = \Delta V_{\text{lin}}^{\text{dist}} \left(1 + \frac{g_i F_3(l_i)}{C_2(l,L)}\right)^{-1}. \tag{S24}$$

Eqs. (S21) and (S24) show that the deviation between the gradient-based estimate and the exact voltage change is controlled by the dimensionless loading terms $g_iF_1/C_1$ and $g_iF_3/C_2$, respectively. The first-order approximation is therefore accurate whenever inhibition acts as a small perturbation to the total effective conductance load. Under this condition, the voltage gradient provides not only a tractable analytical measure of inhibitory efficacy, but also an accurate predictor of the actual finite voltage suppression across inhibitory locations.

### 2. Computational Complexity of Optimal Dendritic Inhibition

We can prove the ODI problem to be NP-complete by Karp reduction.[4] Let M be a deterministic multi-compartment neuron simulated on a discrete horizon T = {1, ..., T} with rational parameters; dendritic space is discretized into locations L. Given excitatory drive ($E: T \to \mathbb{R}^{|L|}$) and candidate inhibitory pulses $P \subseteq L \times T$, for any $S \subseteq P$ denote the somatic voltage by

$$V(t;S) = \text{Sim}(M,E,S,t), \quad t \in \mathcal{T}, \tag{S25}$$

and the no-spike predicate by

$$\text{NoAP}(S) := \left(\max_{t\in\mathcal{T}} V(t;S) < \theta\right). \tag{S26}$$

Decision and optimization forms:

$$\begin{gathered} \text{ODI-DEC: } \exists S \subseteq P,\ |S| \le K \text{ s.t. NoAP}(S)? \\ \text{OPT-ODI: } \min_{S\subseteq P} |S| \text{ s.t. NoAP}(S). \end{gathered} \tag{S27}$$

Verification is one forward simulation over T, hence ODI-DEC ∈ NP. For NP-hardness we work within a monotone-inhibition subclass I_mono:

$$S_1 \subseteq S_2 \Rightarrow V(t;S_2) \le V(t;S_1) \quad (\forall t \in \mathcal{T}). \tag{S28}$$

Reduce Set-Cover (universe U = {$u_1$, ..., $u_n$}, family C = {$C_1$, ..., $C_m$}, budget K) to ODI-DEC. Construct in polynomial time (M, E, P, K) as follows. Partition T into disjoint windows W(u), u ∈ U, with recovery gaps, and set

$$\max_{t\in W(u)} V(t;\varnothing) \geq \theta, \quad \max_{t\notin W(u)} V(t;\varnothing) < \theta, \tag{S29}$$

so each W(u) is an independent thresholding event. Create one pulse $p_j$ ∈ P per set $C_j$ and choose its spatial/temporal profile so that its effectiveness set equals $C_j$:

$$\begin{aligned} u \in C_j &\Rightarrow \max_{t\in W(u)} V\big(t;\{p_j\}\big) < \theta, \\ u \notin C_j &\Rightarrow \max_{t\in W(u)} V\big(t;\{p_j\}\big) \geq \theta. \end{aligned} \tag{S30}$$

By monotonicity,

$$u \in \bigcup_{p_j\in S} C_j \Rightarrow \max_{t\in W(u)} V(t;S) < \theta. \tag{S31}$$

Correctness:

$$\begin{aligned} &\big(\exists C_{i_1}, \dots, C_{i_K} \text{ cover } U\big) \\ \Leftrightarrow &\Big(\exists S = \{p_{i_1}, \dots, p_{i_K}\}, \text{NoAP}(S)\Big). \end{aligned} \tag{S32}$$

The construction uses n windows, m pulses, and T = O(n + m) with polynomial bit-length, hence

$$\text{Set-Cover} \leq_p \text{ODI-DEC}. \tag{S33}$$

Thus ODI-DEC is NP-complete and OPT-ODI is NP-hard.

### 3. Derivation of perturbation kinetics and the compact Φ-a law

The compact formula used in the main text,

$$\delta V_s = \langle \boldsymbol{\Phi}_s, \boldsymbol{a}_{\text{syn}} \rangle, \tag{S34}$$

describes the somatic effect of a single candidate synaptic perturbation around the current background state. Here $\Phi_s$ denotes a state-dependent somatic susceptibility kernel, whereas a_syn denotes the effective local synaptic perturbation. In the main text, the same relationship may be written with the visually simpler shorthand a_syn; in this note, boldface is used whenever the local perturbation is represented as a temporal vector. The dependence on the candidate synaptic event, the readout time, and the current trajectory is suppressed in Eq. (S34) for clarity. Below, we make these dependencies explicit and derive Eq. (S34) from the implicit voltage update implemented in NEURON.[5]

**Implicit compartmental update in NEURON**

Let

$$V(t) = \left[V_1^{(t)}\ V_2^{(t)}\ \cdots\ V_n^{(t)}\right]^{\mathsf{T}} \in \mathbb{R}^n$$

denote the membrane-voltage vector over all physical compartments at discrete time step t, where n is the number of physical compartments.

For a physical compartment i, the backward-Euler discretization of the cable equation used by NEURON can be written as

$$\frac{c_i}{\Delta t}\left(V_i^{(t+1)} - V_i^{(t)}\right) = -I_{m,i}^{(t+1)} + \sum_{k\in N_i} \frac{V_k^{(t+1)} - V_i^{(t+1)}}{r_{ik}}, \tag{S35}$$

where $c_i$ is the compartmental membrane capacitance, Δt is the simulation time step, $N_i$ is the set of neighboring nodes, and r_{ik} is the axial resistance between nodes i and k.

We decompose the transmembrane current into passive leak and all remaining components:

$$I_{m,i}^{(t)} = g_{l,i}\left(V_i^{(t)} - E_{l,i}\right) + I_{\Sigma,i}^{(t)}, \tag{S36}$$

where $g_{l,i}$ and $E_{l,i}$ are the leak conductance and leak reversal potential, and $I_{\Sigma,i}^{(t)}$ includes the non-passive ionic and synaptic currents.

To obtain an explicit linear system for the next-step voltage, NEURON linearizes the non-passive current around the current voltage state:

$$I_{\Sigma,i}^{(t+1)} \approx I_{\Sigma,i}^{(t)} + j_{\Sigma,i}^{(t)}\left(V_i^{(t+1)} - V_i^{(t)}\right), \tag{S37}$$

where

$$j_{\Sigma,i}^{(t)} = \left.\frac{\partial I_{\Sigma,i}}{\partial V_i}\right|_t. \tag{S38}$$

Substituting Eqs. (S36)-(S38) into Eq. (S35) yields, for each physical compartment, a linear relation between the voltages at time steps t + 1 and t.

**Dummy nodes and reduced matrix form**

To enforce current conservation at branch points and terminals, NEURON introduces dummy nodes. These auxiliary nodes have zero membrane capacitance and no transmembrane current, and satisfy

$$\sum_{k\in N_i} \frac{V_k^{(t+1)} - V_i^{(t+1)}}{r_{ik}} = 0 \quad \text{for dummy nodes } i. \tag{S39}$$

Let V_b(t) ∈ R^{n+n_d} denote the voltage vector over both physical and dummy nodes, where n_d is the number of dummy nodes. Collecting the compartment equations and branch constraints gives a linear system of the form

$$A_{\text{full}} V_b^{(t+1)} = B_b^{(t)} V(t) + c_b^{(t)}, \tag{S40}$$

where A_full is determined by passive morphology, axial coupling, capacitance, and branch constraints, while the time dependence enters through the linearized non-passive currents.

We next eliminate the dummy-node degrees of freedom. Let

$$M = SA_{\text{full}}^{-1}, \tag{S41}$$

where S = [I_n 0] selects the physical compartments from the full node vector. Define

$$D = \operatorname{diag}\left(\frac{\Delta t}{c_1}, \dots, \frac{\Delta t}{c_n}\right), \quad J_\Sigma^{(t)} = \operatorname{diag}\left(j_{\Sigma,1}^{(t)}, \dots, j_{\Sigma,n}^{(t)}\right),$$
$$G_l = \begin{bmatrix} g_{l,1} & \cdots & g_{l,n} \end{bmatrix}^\top, \quad E_l = \begin{bmatrix} E_{l,1} & \cdots & E_{l,n} \end{bmatrix}^\top,$$
$$I_\Sigma^{(t)} = \begin{bmatrix} I_{\Sigma,1}^{(t)} & \cdots & I_{\Sigma,n}^{(t)} \end{bmatrix}^\top.$$

With these definitions, the voltage update over the physical compartments can be written as

$$\begin{gathered}\left(I + MDJ_\Sigma^{(t)}\right) V(t+1) = M \\ \left(\left(I + DJ_\Sigma^{(t)}\right) V^{(t)} + D\left(G_l \circ E_l - I_\Sigma^{(t)}\right)\right),\end{gathered} \tag{S42}$$

where ∘ denotes elementwise multiplication.

Define

$$G(t) = I + MDJ_\Sigma^{(t)}. \tag{S43}$$

Assuming G(t) is invertible, Eq. (S42) yields

$$V^{(t+1)} = A^{(t)} V^{(t)} + \eta^{(t)}, \tag{S44}$$

with

$$A^{(t)} = \left(G^{(t)}\right)^{-1} M \left(I + DJ_\Sigma^{(t)}\right), \tag{S45}$$

and

$$\eta^{(t)} = \left(G^{(t)}\right)^{-1} MD\left(G_l \circ E_l - I_\Sigma^{(t)}\right). \tag{S46}$$

Equation (S44) is the compact voltage recursion used in the main text. Importantly, both $A^{(t)}$ and η(t) are evaluated on the current trajectory and therefore depend implicitly on the current voltage state through $J_\Sigma^{(t)}$ and $I_\Sigma^{(t)}$.

**One-step voltage map and its Jacobian**

To make this dependence explicit, we define the one-step voltage map

$$V^{(t+1)} = F^{(t)}\left(V^{(t)}\right), \tag{S47}$$

with

$$F^{(t)}(V) = A^{(t)}(V)\, V + \eta^{(t)}(V), \tag{S48}$$

where

$$A^{(t)}(V) = \left(G^{(t)}(V)\right)^{-1} M\big(I + DJ_{\Sigma}(V)\big), \tag{S49}$$

and

$$\eta^{(t)}(V) = \left(G^{(t)}(V)\right)^{-1} MD\big(G_l \circ E_l - I_{\Sigma}(V)\big), \tag{S50}$$

with

$$G^{(t)}(V) = I + MDJ_{\Sigma}(V). \tag{S51}$$

Thus, Eq. (S44) is the trajectory-evaluated form of Eq. (S47). Because both $A^{(t)}(V)$ and $\eta^{(t)}(V)$ depend on V, the first-order propagation of perturbations is governed not by $A^{(t)}$ itself, but by the Jacobian of the one-step map:

$$K^{(t)} = \left.\frac{\partial F^{(t)}}{\partial V}\right|_{V=X^{(t)}}, \tag{S52}$$

where $X^{(t)}$ denotes the current background trajectory at time step t.

Equivalently, if Eq. (S48) is expanded,

$$K^{(t)} = A^{(t)} + \left.\frac{\partial A^{(t)}(V)}{\partial V}\right|_{V=X^{(t)}} \lrcorner X^{(t)} + \left.\frac{\partial \eta^{(t)}(V)}{\partial V}\right|_{V=X^{(t)}}, \tag{S53}$$

where the contraction indicates the action of the derivative tensor of $A^{(t)}(V)$ on the current state vector.

**First-order response to a single candidate synaptic perturbation**

We now consider a single candidate synaptic event inserted at a specified dendritic location and time. Let $\epsilon_{\text{syn}}$ denote a formal local differential parameter for this candidate event, and let X denote the current forward trajectory of the background system. All response quantities below are defined with respect to this current trajectory.

We denote the first-order voltage response to the candidate perturbation by

$$\delta V^{(t)} \equiv \frac{\partial V^{(t)}}{\partial \epsilon_{\text{syn}}}.$$

Here $\epsilon_{\text{syn}}$ is used only to derive the local first-order response. In the fixed-strength inhibitory events used for optimal dendritic inhibition, the event strength is not treated as an independently optimized variable; instead, it is absorbed into the effective local perturbation waveform $a_{\text{syn}}^{(t)}$ defined below.

Differentiating Eq. (S47) with respect to $\epsilon_{\mathrm{syn}}$ along X gives

$$\delta V^{(t+1)} = K^{(t)} \delta V^{(t)} + \xi^{(t)}, \tag{S54}$$

where

$$\xi^{(t)} = \left. \frac{\partial F^{(t)}}{\partial \epsilon_{\mathrm{syn}}} \right|_{X^{(t)}} \tag{S55}$$

is the direct perturbation source term induced by the candidate synaptic event at time step t. This notation separates the source term $\xi^{(t)}$ in the perturbation recursion from the affine term $\eta^{(t)}$ in the voltage recursion of Eq. (S44).

**Effective local perturbation waveform**

For a synaptic perturbation localized at a fixed candidate site, the source term can be factored into a spatial injection vector and a scalar temporal waveform:

$$\xi^{(t)} = B_{\mathrm{syn}}^{(t)} a_{\mathrm{syn}}^{(t)}, \tag{S56}$$

where $a_{\mathrm{syn}}^{(t)}$ is the effective local perturbation generated by the candidate synaptic event at time step t, and $B_{\mathrm{syn}}^{(t)}$ specifies how a unit local perturbation enters the full voltage system after the implicit update has been resolved. For a fixed-strength synaptic event, $a_{\mathrm{syn}}^{(t)}$ contains the fixed event strength, the synaptic time course, the local driving force, and the sign of the perturbation. Thus, once the candidate location and timing are specified, $a_{\mathrm{syn}}^{(t)}$ is determined rather than independently optimized.

Substituting Eq. (S56) into Eq. (S54) gives

$$\delta V^{(t+1)} = K^{(t)} \delta V^{(t)} + B_{\mathrm{syn}}^{(t)} a_{\mathrm{syn}}^{(t)}. \tag{S57}$$

This is the perturbation recursion underlying the main-text response law. The first term propagates the previously accumulated response through the local Jacobian dynamics, whereas the second term injects the newly generated local synaptic perturbation at the candidate site.

**Temporal propagation and somatic readout**

Assuming zero initial response,

$$\delta V^{(0)} = 0, \tag{S58}$$

define the transition operator

$$P^{(t,k+1)} = K^{(t-1)} K^{(t-2)} \cdots K^{(k+1)}, \quad P^{(t,t)} = I. \tag{S59}$$

Repeated substitution of Eq. (S57) yields

$$\delta V^{(t^*)} = \sum_{k=0}^{t^*-1} P^{(t^*,k+1)}\, B_{\text{syn}}^{(k)} a_{\text{syn}}^{(k)}, \tag{S60}$$

where t* is the readout time.

Let the somatic voltage be

$$V_s^{(t)} = e_s^{\top} V(t), \tag{S61}$$

where $e_s$ is the basis vector selecting the soma. Projecting Eq. (S60) onto the soma gives

$$\delta V_s^{(t^*|X)} = \sum_{k=0}^{t^*-1} e_s^{\top}\, P^{(t^*,k+1)} B_{\text{syn}}^{(k)} a_{\text{syn}}^{(k)}. \tag{S62}$$

We therefore define the site-to-soma susceptibility kernel

$$\Phi_s^{(t^*,k|X)} = e_s^{\top} P^{(t^*,k+1)} B_{\text{syn}}^{(k)}, \quad k = 0,1,\dots,t^*-1. \tag{S63}$$

Equation (S62) then becomes

$$\delta V_s^{(t^*|X)} = \sum_{k=0}^{t^*-1} \Phi_s^{(t^*,k|X)}\, a_{\text{syn}}^{(k)}. \tag{S64}$$

**Compact notation used in the main text**

For a fixed candidate synaptic event, a fixed readout time t*, and a fixed current trajectory X, Eq. (S64) is a temporal inner product between a susceptibility vector and a local perturbation vector. Define

$$\Phi_s = \left[\Phi_s^{(t^*,0|X)}\ \Phi_s^{(t^*,1|X)}\ \cdots\ \Phi_s^{(t^*,t^*-1|X)}\right]^{\top}, \tag{S65}$$

and

$$a_{\text{syn}} = \left[a_{\text{syn}}^{(0)}\ a_{\text{syn}}^{(1)}\ \cdots\ a_{\text{syn}}^{(t^*-1)}\right]^{\top}. \tag{S66}$$

Then Eq. (S64) can be written as

$$\delta V_s^{(t^*|X)} = \langle \Phi_s, a_{\text{syn}} \rangle. \tag{S67}$$

Suppressing the dependence on the readout time t*, the current trajectory X, and the candidate synaptic event gives the compact Φ-a law used in the main text:

$$\delta V_s = \langle \boldsymbol{\Phi}_s, \boldsymbol{a}_{\text{syn}} \rangle. \tag{S68}$$

In this compact notation, $\Phi_s$ represents the state-dependent global susceptibility from the candidate synaptic perturbation to the soma, whereas a_syn represents the effective local synaptic perturbation. Because the dendritic system is nonlinear, the susceptibility vector depends on the current trajectory and is recomputed after each accepted inhibitory perturbation. Therefore, Eq. (S68) describes a local first-order effect around the current state and does not imply linear superposition across multiple synaptic inputs.

### Supplementary References


1 Rall, W. Branching dendritic trees and motoneuron membrane resistivity. Exp Neurol 1, 491-527 (1959).

2 Koch, C., Poggio, T. & Torre, V. Nonlinear interactions in a dendritic tree: localization, timing, and role in information processing. Proc Natl Acad Sci USA 80, 2799-2802, doi:10.1073/pnas.80.9.2799 (1983).

3 Gidon, A. & Segev, I. Principles governing the operation of synaptic inhibition in dendrites. Neuron 75, 330-341, doi:10.1016/j.neuron.2012.05.015 (2012).

4 Karp, R. M. Reducibility among combinatorial problems. In Complexity of Computer Computations (eds Miller, R. E. & Thatcher, J. W.) 85-103 (Plenum, 1972).

5 Hines, M. L. & Carnevale, N. T. The NEURON simulation environment. Neural Comput 9, 1179-1209, doi:10.1162/neco.1997.9.6.1179 (1997).

### Supplementary Figures

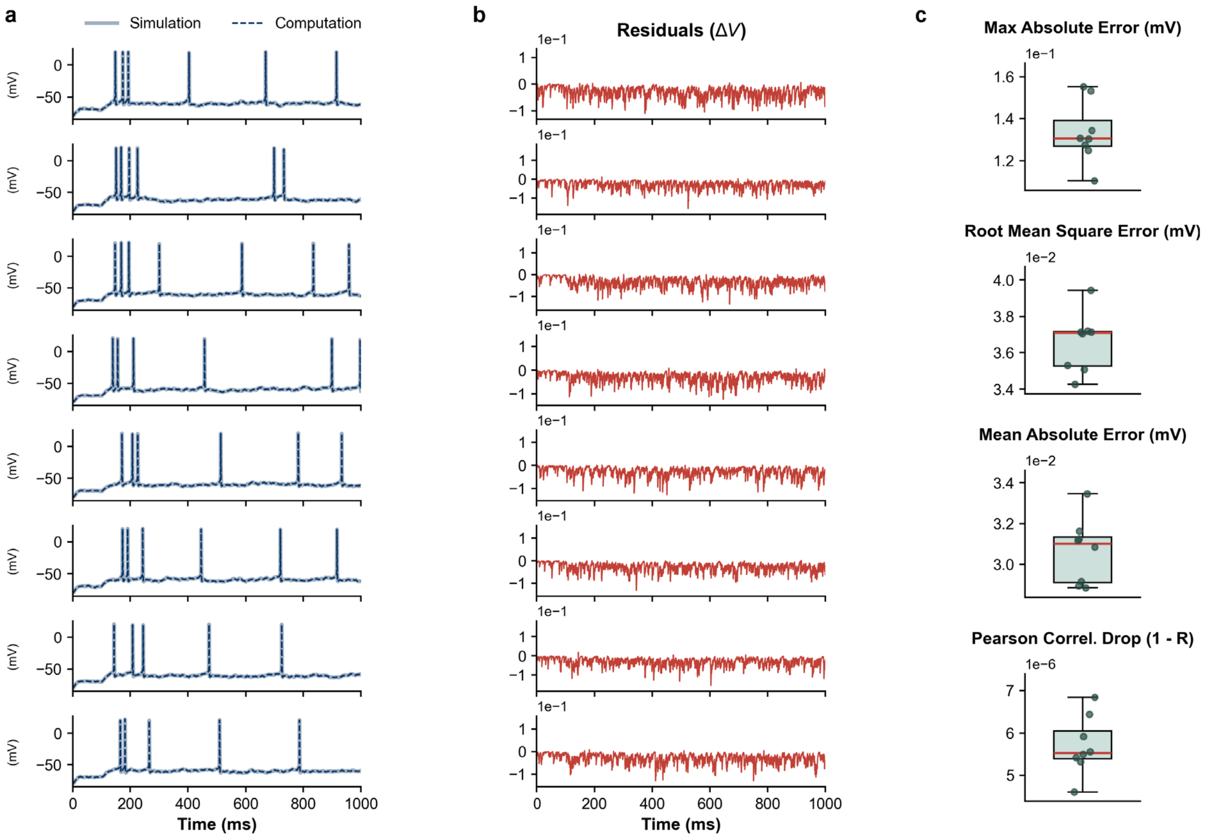


**Supplementary Fig. 1 | Matrix recursion reproduces NEURON trajectories in detailed simulations.** a, Representative somatic voltage traces generated by the original NEURON simulation and by the reformulated matrix computation. b, Residual voltage differences (simulation minus computation) for the corresponding traces. c, Summary error metrics across simulations: maximum absolute error, root mean square error, mean absolute error and Pearson correlation drop (1 − R). Errors remain near numerical tolerance, validating the simulator-level reformulation.

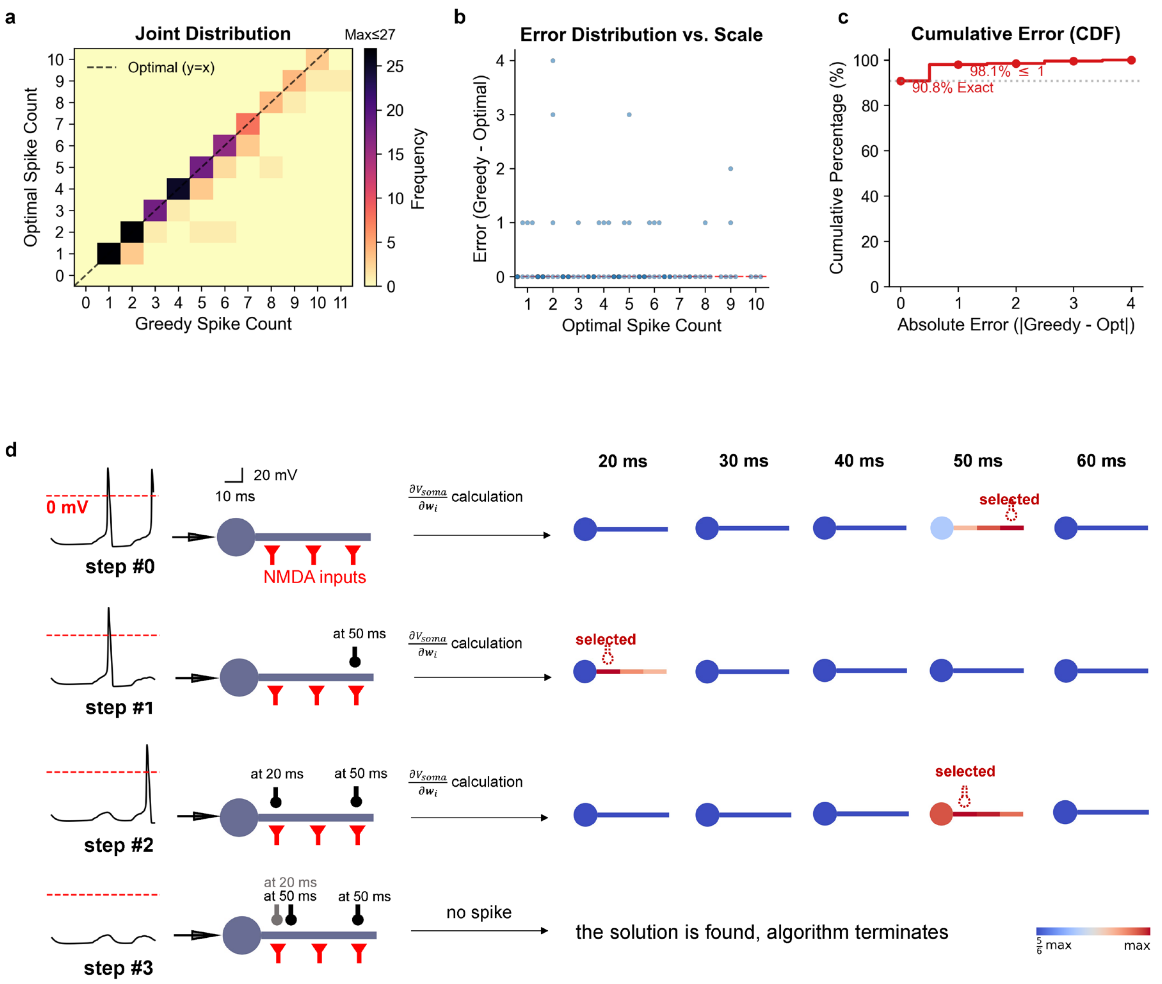


**Supplementary Fig. 2 | Greedy ODI solutions closely match exhaustive optima in a toy neuron.** a, Joint distribution of inhibitory spike counts returned by the greedy algorithm and by exhaustive search; dashed line indicates equality. b, Error in spike count versus the exact optimum. c, Cumulative distribution of absolute error, showing exact or near-exact solutions for most cases. d, Example greedy search trajectory in a ball-and-stick model. Starting from NMDA-evoked somatic spiking, the algorithm repeatedly computes $\delta V_s := \partial \mathrm{V_{soma}} / \partial \mathrm{w_i}$ over candidate inhibitory pulses, selects the strongest vetoing pulse, updates the trajectory and terminates when somatic spiking is suppressed.

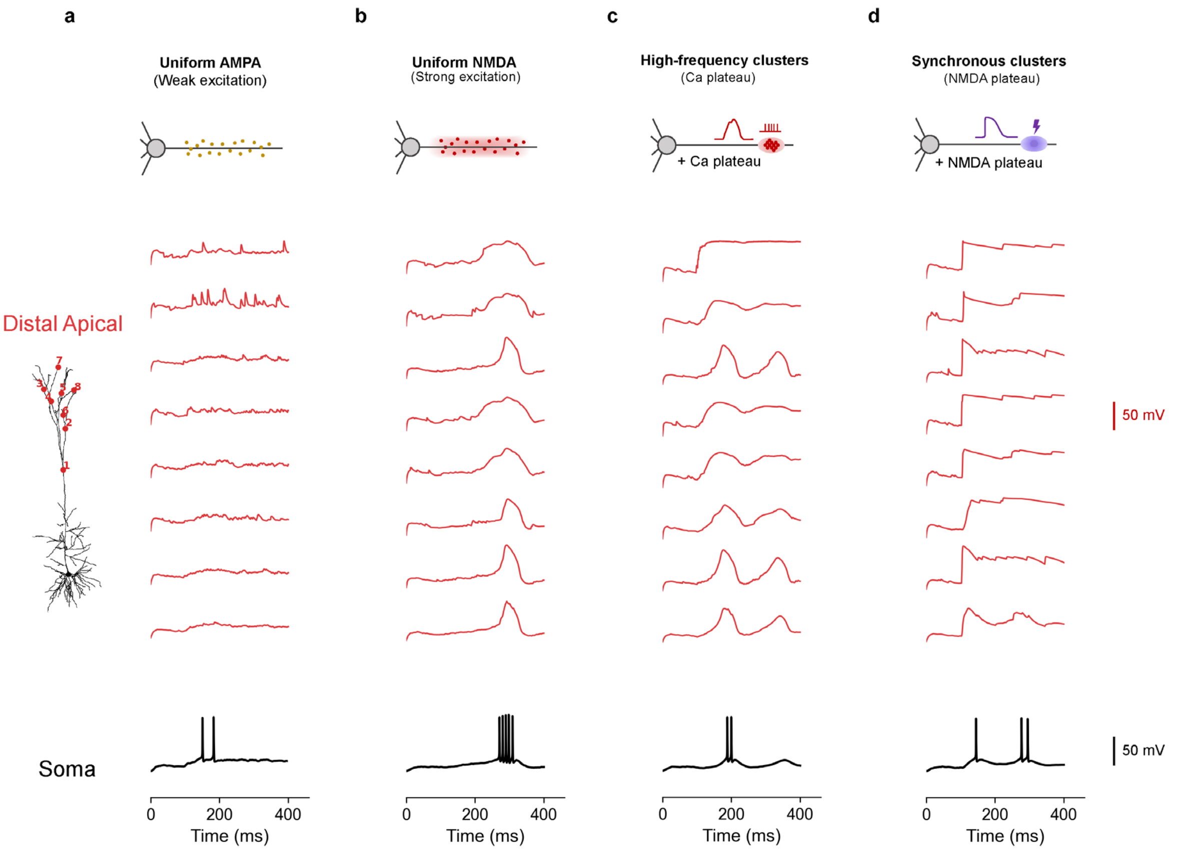


**Supplementary Fig. 3 | Excitation paradigms used to recruit distinct dendritic states in the L5 model.** a–d, Schematic input patterns, distal dendritic voltage traces and somatic voltage responses for uniform AMPA (a), uniform NMDA/AMPA (b), high-frequency distal NMDA/AMPA clusters (c) and synchronous distal NMDA/AMPA clusters (d). Red traces show voltage responses at numbered distal apical sites; black traces show somatic voltage. Uniform AMPA evokes weak distributed activity, uniform NMDA produces stronger distributed depolarization, high-frequency clusters evoke $Ca^{2+}$ plateau-like events, and synchronous clusters evoke NMDA plateau-like events. Scale bars, 50 mV.

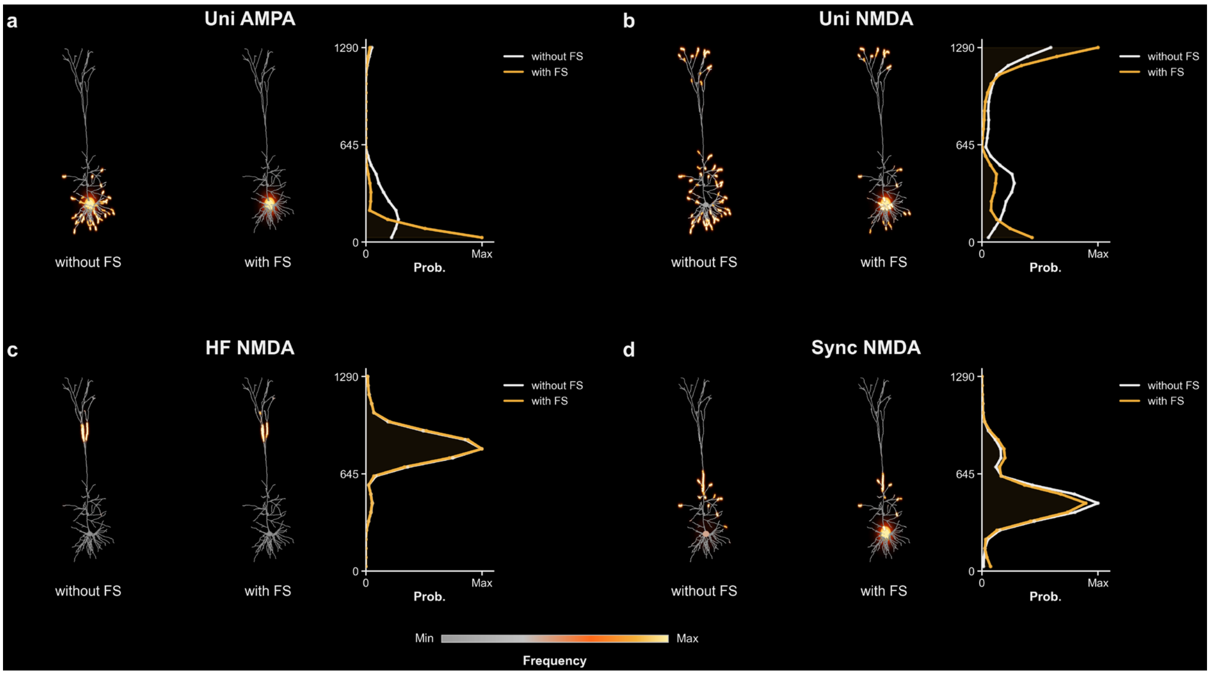


**Supplementary Fig. 4 | Fast-spiking perisomatic candidates primarily affect inhibition under unstructured excitation.** ODI maps and radial probability profiles computed with and without fast-spiking (FS) perisomatic inhibitory candidates for uniform AMPA, uniform NMDA/AMPA, high-frequency distal NMDA/AMPA and synchronous distal NMDA/AMPA excitation. Adding FS candidates near the soma increases the perisomatic/proximal component of optimal inhibition under unstructured AMPA or NMDA excitation, but has little effect on structured distal regimes, in which inhibition remains concentrated near the distal source or intermediate apical pathway. Warm colours denote higher selection probability; profiles show selection probability as a function of distance from the soma.
